# Molecular signatures of resource competition: clonal interference favors ecological diversification and can lead to incipient speciation

**DOI:** 10.1101/2020.11.20.391151

**Authors:** Massimo Amicone, Isabel Gordo

## Abstract

Microbial ecosystems harbor an astonishing diversity that can persist for long times. To understand how such diversity is structured and maintained, ecological and evolutionary processes need to be integrated at similar timescales. Here, we study a model of resource competition that allows for evolution *via de novo* mutation, and focus on rapidly adapting asexual populations with large mutational inputs, as typical of many bacteria species. We characterize the adaptation and diversification of an initially maladapted population and show how the eco-evolutionary dynamics are shaped by the interaction between simultaneously emerging lineages – clonal interference. We find that in large populations, more intense clonal interference fosters diversification under sympatry, increasing the probability that phenotypically and genetically distinct clusters stably coexist. In smaller populations, the accumulation of deleterious and compensatory mutations can push further the diversification process and kick-start speciation. Our findings have implications beyond microbial populations, providing novel insights about the interplay between ecology and evolution in clonal populations.

## Introduction

Understanding the mechanisms behind the evolution of biodiversity and the formation of communities remains a difficult challenge. One must integrate ecology and evolution over similar timescales, as taken together, they can give rise to phenomena that could not be explained by either alone (Schoener, 2011). The competitive exclusion principle (first stated by Hardin, 1960) theoretically binds the number of species by the number of limiting resources. This principle generated an apparent contradiction between theoretical expectations and observations, often referred to as the “paradox of the plankton” (Hutchinson, 1961). In fact, ecosystems can be replete of diversity even in limiting environments, both in nature (Hutchinson, 1961; Tilman, 1982; Huston, 1994) and in more controlled laboratory conditions (Maharjan *et al*., 2006; Gresham *et al*., 2008; Kinnersley, Holben and Rosenzweig, 2009; Herron and Doebeli, 2013; Good *et al*., 2017). Different theoretical approaches have been adopted to resolve such controversy. One approach is to assume an existing diversity and identify mechanisms that can maintain it. Following this, several ecological properties were proposed to maintain diversity, including heterogeneity in space (Abrams, 1988) and time (Litchman and Klausmeier, 2001), trade-offs on the species’ traits (Posfai, Taillefumier and Wingreen, 2017), or gene regulation (Pacciani-Mori *et al*., 2020). Here we explore a classical model of competition for resources, where extensive diversity can be maintained by a metabolic trade-off (Posfai, Taillefumier and Wingreen, 2017), and ask a different question: in an initially monomorphic population, what diversity can be generated and maintained if the species’ traits continuously evolve?

In eco-evolutionary frameworks, mutations generate new genetic variants whose fate depends on the state of the ecosystem and, in turn, their increase in frequency can alter the populations. A common outcome of such eco-evolutionary feedbacks is that evolution limits diversity by reducing the effectiveness of coexistence mechanisms (Edwards *et al*., 2018). The diversity that would be possible by ecological principles alone, is reduced by selection of the fittest and competitive exclusion. Several studies have produced novel understanding on the evolution of diversity (Dieckmann and Doebeli, 1999; Shoresh, Hegreness and Kishony, 2008; Doebeli, 2011; Kremer and Klausmeier, 2017), but the majority rely on the strong-selection-weak-mutation assumption, i.e. on a timescale separation between ecological and evolutionary processes (but see Farahpour *et al*., 2018). The emergence of mutations is assumed to be much slower than the ecological dynamics, thus, before a new lineage arises, the population has already reached ecological equilibrium. While allowing for analytical tractability the weak mutation assumption comes at a cost: it neglects the overlap between multiple evolving lineages – clonal interference. Clonal interference has been extensively observed in microbial communities *in vitro* and *in vivo* (Desai, Fisher and Murray, 2007; Barroso-Batista *et al*., 2014) and can occur under different regimes of intensity: a weak regime where a few lineages compete for fixation (Philip J. Gerrish & Richard E. Lenski, 1998; Billiard and Smadi, 2020) or one where many different haplotypes segregate (Good *et al*., 2012). Population genetics models incorporating clonal interference have generated predictions for the adaptation rate, fixation probabilities and genetic diversity in a population (Gerrish and Lenski, 1998; Park and Krug, 2007; Good *et al*., 2012; de Sousa *et al*., 2016), yet they typically ignore ecological interactions (but see Good, Martis and Hallatschek, 2018).

When the population size *(N)* and/or the mutation rate to new beneficial mutations *(Ub)* are not small *(NUb* >>1), multiple lineages can increase in frequency simultaneously, ecologically interact with each other and evolve in non-trivial ways. Although these processes are inevitably intertwined in real ecosystems (Lawrence *et al*., 2012; Barroso-Batista *et al*., 2014, 2020; Garud *et al*., 2019), theoretical work is still needed to investigate how they act in chorus.

Good and colleagues (Good, Martis and Hallatschek, 2018) have recently developed a theoretical work that incorporates ecological and evolutionary mechanisms *via* a combination of frequency-dependent and directional selection. Their eco-evolutionary model is able to reproduce empirical patterns of co-existence and fixation of new mutations in experimentally evolved clonal populations (Good *et al*., 2017). It also shows how diversification depends on the ratio between the rates of strategy mutations and unconditionally beneficial mutations. However, in order to derive analytical expression for the eco-evolutionary dynamics, the authors have focused on the weak mutation limit *NU_b_* <<1 and only briefly investigated clonal interference.

Here, we study a similar model to that of (Posfai, Taillefumier and Wingreen, 2017; Good, Martis and Hallatschek, 2018) but assume a trade-off that only affects well adapted genotypes (fitness-dependent trade-off) and conduct a more systematic simulation study of the different mutation regimes, including extensive clonal interference, in large and small populations. We follow an initially isogenic population throughout time and characterize the patterns of adaptation at both phenotypic and genetic levels, by common statistics used to analyze molecular evolution data. We focused only on mutations that affect the ability of consuming resources (instead of differentiating between strategy and unconditionally beneficial mutations) and we do not impose restrictive assumptions on mutation rates and on timescales, as common in other models (Geritz *et al*., 1998; Shoresh, Hegreness and Kishony, 2008; Good, Martis and Hallatschek, 2018). Albeit at the cost of analytical tractability, our approach is to focus on the phenomena that emerge from the stochastic simulations where many more lineages compete, compared to previous studies (Billiard and Smadi, 2020).

We find that: high levels of intra-specific variation can be generated and maintained via a balance between selection and mutation; functionally distinct clusters of genotypes – ecotypes – can emerge and stably coexist; and the interaction between large mutational inputs and the energetic trade-off can lead to incipient speciation. Taken together, our results describe how clonal populations can give rise to extensive diversity and establish a first form of community, even in simple and constant environments.

## Model and Methods

### Eco-Evolutionary model

We model the dynamics of a single clonal lineage evolving to consume a set of different substitutable resources, constantly replenished in a well-mixed environment (Fig. 1A). Individuals mutate at a rate *U* (per-genome, per-generation rate of non-lethal mutations) and the fate of the emerging mutations depends on their phenotypic effects, on the resource concentration, on the other individuals present in the environment and on drift.

**Figure 1.**
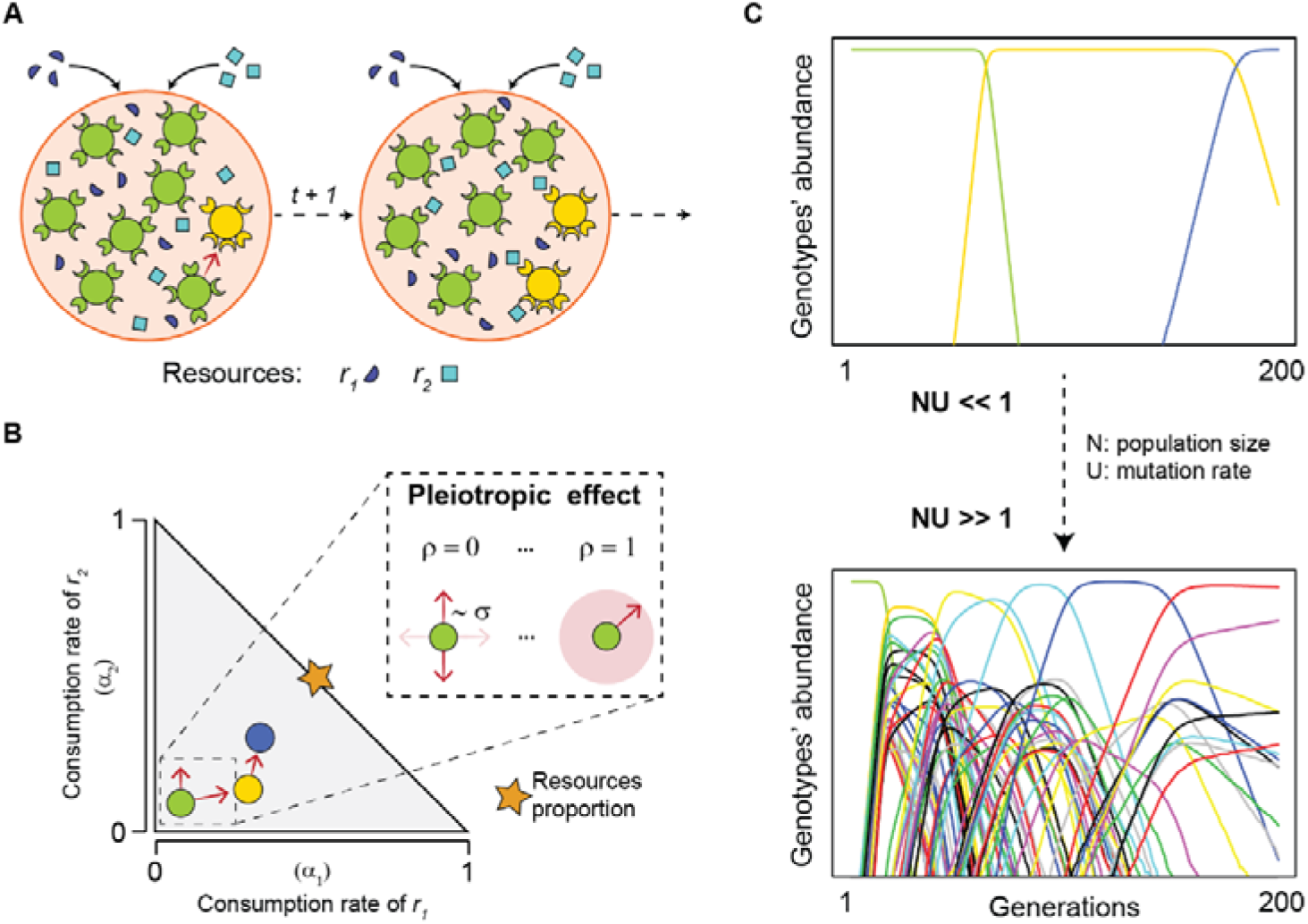
Ecological dynamics and individual-based evolutionary processes. **A)** Illustration of the eco-evolutionary dynamics. Bacterial genotypes, represented by circles of different colors, grow according to the constant input of resources (squares and semicircles) and their phenotypic traits, represented by the enzyme-like structures on the circles. Mutation events (red arrow) can generate new types whose fate will depend on drift and selection. **B)** Constrained phenotypic space and mutation process. An initially maladapted monomorphic population (green circle) can acquire de-novo mutations according to the given assumptions (as explained in the inset) and move inside the phenotypic space with an upper bound on the total energy. **C)** Examples of different mutation regimes: from low/absent (NU << 1) to extensive (NU >>1) clonal interference. Each color represents a different genotype.

The underlying dynamics are based on the MacArthur’s consumer resource model (Mac Arthur, 1969), recently formalized to explain high levels of diversity in the presence of a metabolic trade-off (Posfai, Taillefumier and Wingreen, 2017) and further extended to study adaptation (Good, Martis and Hallatschek, 2018). Briefly, let *M* be the number of types present at time *t* with densities (#*cells/V*) *n_i_*, (*i*=1 …*M*) and *R* the number of substitutable resources with input concentrations *r_j_*, {*j*=1…*R*). The expected density dynamics of each type are:

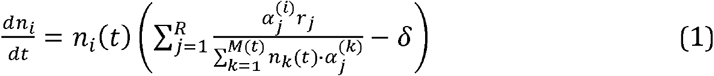

where 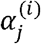 represents the consumption rate of resource *j* by type *i* and δ is the death rate. The resource amounts are constant in this model since, as *Posfai et al,* we assume that metabolic reactions occur much faster than cell division (Posfai, Taillefumier and Wingreen, 2017).

We assume a finite amount of energy available for each cell and limit their ability of consuming resources by an energetic constraint (*E*):

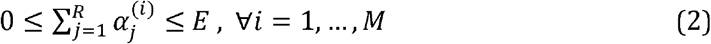

Under this assumption, *E* acts as an upper bound and not as a fixed energy budget, as previously investigated (Posfai, Taillefumier and Wingreen, 2017; de Oliveira, Amado and Campos, 2018; Amado and Campos, 2019). Assuming equally supplied resources (*r_j_* = *r* ∀*j*) and unitary energy, volume and death rate [*E, V δ* =1}, the population size is *N=Rr*.

We model an initial isogenic population (M(t_0_) =1) with given traits 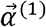 and allow for mutations that change the heritable traits and give rise to new genotypes. Every generation, each genotype *i* (*i=1…M*(*t*)) will generate a Poisson-distributed number of mutants with expected value *n_i_*(*t*) · *U*. Assuming an infinite site model, a mutation on genotype *i* will result into a new individual with unique genotype (*i*’) whose phenotypes differ from the parental traits by a small amount: 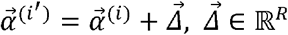.

The mutation effects are drawn from a normal distribution as follows:

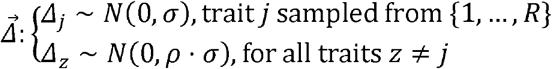

If *ρ* =1, a mutation changes all the traits with equal probability. If 0 < *ρ* <1, mutations target one trait (randomly sampled with probability 1/*R*), but partially alter the other ones. If *ρ* =0, a mutation only changes a single trait. The parameter *ρ* modulates different degrees of trait interdependence or equivalently the pleiotropic effect of mutations, while the parameter *σ* modulates the magnitude of the mutation effects (see Fig. 1B).

In order to respect the boundary condition (2), we assumed that: i) mutations leading to negative values of *α_j_* are loss of function and thus assigned *α_j_*=0; ii) mutations that do not respect the energy constraint cannot exist, therefore Δ*j* are drawn until the upper limit of (2) is satisfied.

In the limit of discrete time steps, we define the selection acting on genotype *i* at time *t*, *S_i_*(*t*), as the expected increase in abundance in the absence of drift, such that *E*[*n_i_*(*t* + 1)] = *n_i_*(*t*)(1 + *S_i_*(*t*)), and from (1):

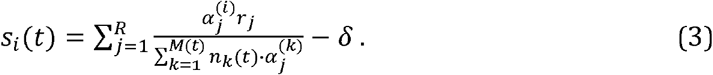

The fate of each genotype depends on its ability to consume each of the resources and on the ecosystem’s ecology.

To simulate drift, we draw the final abundances via multinomial sampling with probabilities 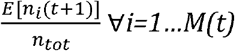. Every generation, the number of genotypes *M*(*t*+1) is updated together with the relative abundances, traits and phylogeny.

### Numerical simulations

In each simulation, an initially maladapted 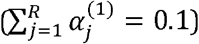 and monomorphic population undergoes the eco-evolutionary process described above for *10^4^* generations, enough to reach phenotypic equilibrium. We focus on the role of different mutation types and regimes (Fig. 1B-C), thus, the main results are obtained via exploring different ranges of the parameters *N, U, σ* and *ρ*, whose values are specified along the text. Each combination of parameters was simulated in 100 independent replicates to obtain the statistics of diversity. In order to disentangle the role of the energetic constraint assumption, we also simulated adaptation under selection but without any boundary condition (or equivalently E=+∞ for (2)). The algorithm was written in *R (version 3.6.1)* and the results analyzed in *RStudio. We* validated the code by comparing simulations outcomes against well-known theoretical expectations from population genetics (see Fig. S1). The code for the simulations is available at https://figshare.com/s/a85bfd4b9f64afaebalb.

### Neutral mutation model

Neutral theories provide null expectations for the genetic diversity within a population, assuming that this population has reached equilibrium. As the time to reach such equilibrium is proportional to the population size (N), neutral predictions are not adequate for “short-term” adaptation of large populations *(T* << *N*), e.g. experimental evolution with bacterial populations. Thus, we run simulations with only drift, for the same time as the selection case (*T* = 10^4^). These out-of-equilibrium simulations help disentangling the contribution of neutral processes to the patterns that we observe during adaptation under selection.

Under the neutral mutation model, genotypes acquire mutations with the same trait effects and probability as described before, but their growth probabilities are equal and do not depend on the phenotypes. Modelling the explicit *α*_*j*_ under neutrality, instead of assuming that the mutations have no effect (*Δ_j_*=0), allows for both genetic and phenotypic comparisons with the model of selection.

### Phenotypic and genotypic diversity

Within a population, each type *i* is characterized by a vector of its consumption traits 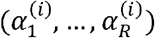 and a vector of the mutations that gave rise to it, each with a unique identifier (e.g. the vector [1,2,7,10] represents genotype 10 whose ancestors are, in order, genotypes 7,2 and 1, and 1 is the ancestor common to every type). From this implementation we can reconstruct the entire phylogeny of a population at any time point and map it on the phenotypic space.

We measure the genetic diversity of a population by the average pairwise genetic distance *π_G_* in a sample of *m* individuals: 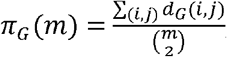, where *m=100* and *d_G_*(*i,j*) is the number of mutations that separate genotype *i* from *j*. From *π_G_* and the number of segregating sites in the sample, we further compute another population genetics statistic: Tajima’s D (Tajima, 1989). The expected value of D at equilibrium is: zero, without selection; positive, under divergent selection; and negative, under purifying selection. However, as the time of our focus (10^4^ generations) is much shorter than that required to achieve equilibrium (~*N*=10^7^), only relative comparisons are meaningful. At the functional level, we compute the average pairwise phenotypic distance *π_P_*, defined as: 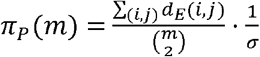, where *d_E_* is the classical Euclidean distance and it is normalized *σ*, for a direct comparison with *π_G_*. For each of the evolved populations, we identified functional clusters from their phenotypic distribution, via the mean shift clustering algorithm (Cheng, 1995), implemented through the *R* package *meanShiftR* version 0.53 (Lisic, 2018) (see Appendix S2 in Supporting Information).

## Results

### Competition-driven diminishing return and the rate of adaptation

The initially monomorphic population, which is poorly adapted, is expected to acquire mutations that improve the ability to consume the available resources and to advance in the phenotypic space towards better adapted states. Our first aim is to identify what influences the speed of this adaptive process.

Under the competition for resources set by (1), how does selection change over time and with the genetic composition of the population? Selection acting on an emerging genotype *i* is given by (3) and depends on the population investment on each resource 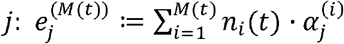. Let us first simplify the problem by considering a monomorphic (*M=1*) population whose phenotypes 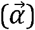 mirror the resource input proportions: 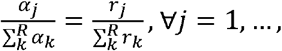 which consists of the local optimal strategy for a given energetic investment 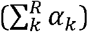. In the absence of mutations, such population at equilibrium satisfies:

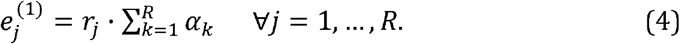

Now consider a mutant that emerges from this population with phenotypes 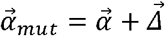. From (3) and (4) it follows that the selection acting on such mutant is:

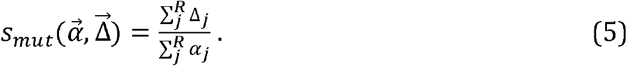

While bigger steps result in stronger selection, equation (5) also implies that the same mutations are subject to weaker selection when emerging on a better adapted background. Thus, competition-driven selection in our system, as in (Good, Martis and Hallatschek, 2018), exhibits diminishing return epistasis – the benefit decline in populations with higher mean fitness – consistent with many empirical observations in microbial populations (Chou *et al*., 2011; Kryazhimskiy *et al*., 2014; Schoustra *et al*., 2016; Wünsche *et al.*, 2017).

Because we assume phenotypic changes follow a normal distribution, their additive effect will also follow a normal distribution: 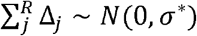 where 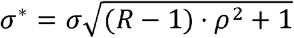; this implies that stronger pleiotropic effects lead to stronger selection. From the continuous univariate distribution theory (Johnson *et al*., 1994) we can retrieve the expected beneficial mutation effect 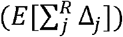 and, from (5), compute the corresponding expected selection coefficients (*E*[*s*^+^]) for varying values of 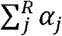 (see Appendix S1 in Supporting Information). Figure 2A shows how the strength of positive selection decreases for better adapted genetic backgrounds across different *σ* and *ρ* conditions. It is important to note that the diminishing return epistasis in this model is not due to the energy constraint; nonetheless, such boundary condition makes the deleterious mutations more common (see Fig. S2 and Appendix S1) and further slows down the rate of adaptation of these well adapted populations (inset in Fig. 2A).

**Figure 2.**
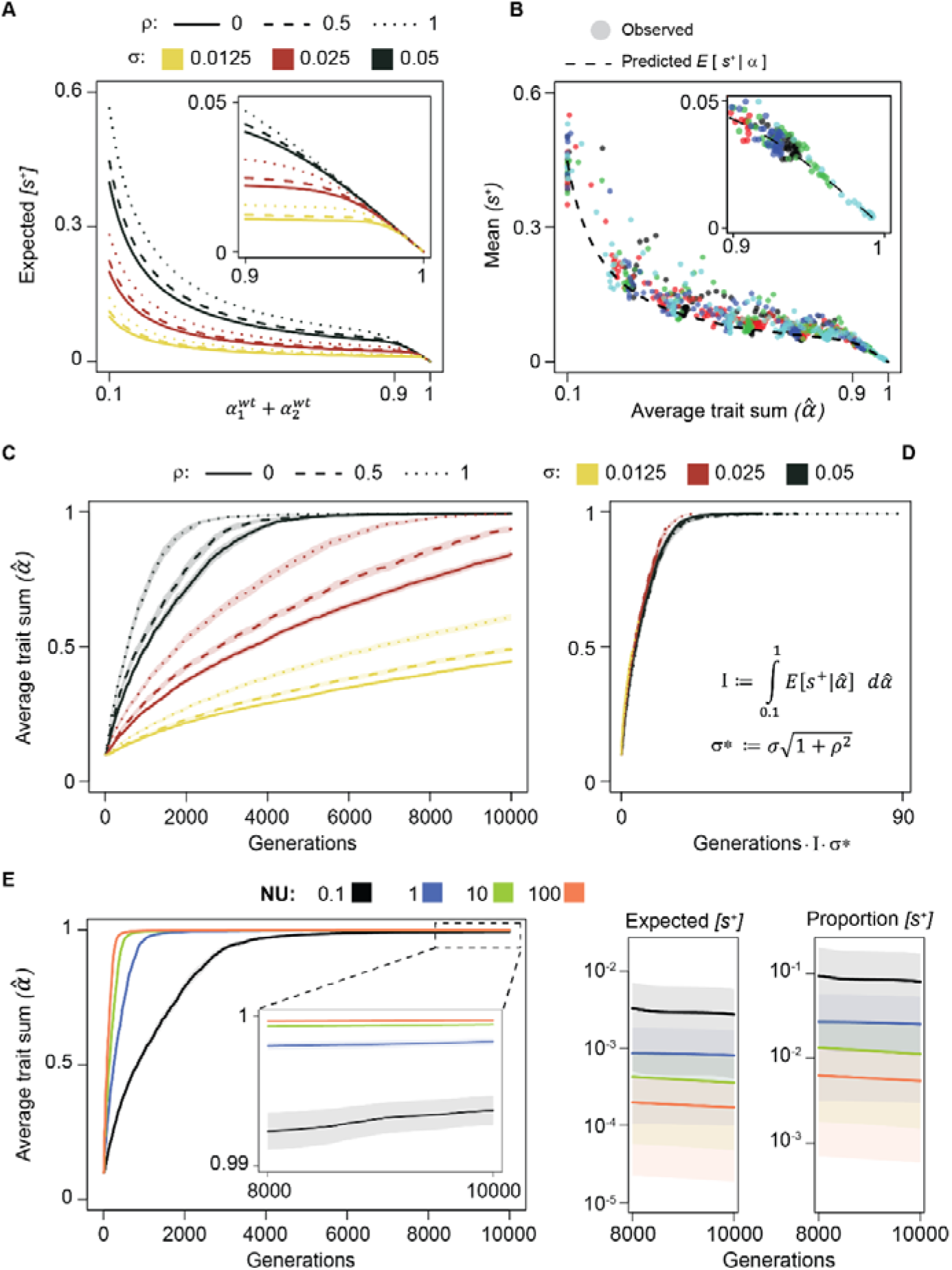
Diminishing return epistasis and the adaptation rate. **A)** Analytical predictions of the diminishing return epistasis in monomorphic populations. The inset shows the effect of the energy constraint **B)** Dots represent the mean of the observed positive selection during the first 300 generations of adaptation. The dotted line represents the prediction using the average population trait sum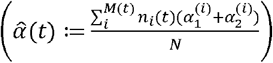. Each color represents an independent simulated population, which adapted with σ = 0.05, ρ = 0.5, *N* = 10^7^ and *U*= 10^-5^. **C)** The population average trait sum 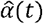 is shown as proxy of adaptation under different *σ* and *ρ* conditions. Other parameters: *R* = 2, *N* = 10^7^, *U* = 10^-8^. Lines are the averages over 100 simulations and the shaded areas represent the confidence interval. **D)** Same dynamics as in C, but on a different scale, as defined in the figure. E) Phenotypic adaptation across different mutational inputs (left) and expected strength and proportion of beneficial mutations at the end ofthe adaptation process. Other parameters for panel E: *N* = 10^7^, *σ* = 0.05, p = 0.5.

Next, we tested how well the approximation obtained by assuming monomorphic populations at consecutive equilibria, predicts what happens in regimes where the adapting populations are polymorphic and out of equilibrium. From the simulations with *NU=100* and two resources, we computed the average population trait sum 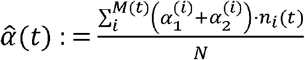 and the expected beneficial selection coefficient as 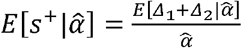. We then compare this expectation with the mean beneficial selection observed in the simulations, during the first 300 generations. The strength of selection acting on polymorphic populations follows the predicted diminishing return pattern, but is often underestimated (Fig. 2B). In fact, it decreases with the mean population phenotype 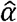, but it increases with the population phenotype variance (Fig. S3).

Numerical simulations further allow us to link the mutation types with the speed of phenotypic adaptation: larger phenotypic changes imply stronger selection, resulting in faster adaptation (Fig. 2C). We find that, when time is scaled by 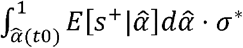, the populations’ mean phenotype moves with similar velocity, demonstrating that both the complex form of selection and the mutation type mediate the speed of adaptation (Fig. 2D and Appendix S1).

Finally, the simulations show that, as expected, larger mutational inputs accelerate phenotypic adaptation and more rapidly lead the populations to a quasi-neutral regime where beneficial mutations are rare and of small effects 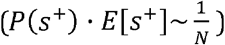 (Fig. 2E).

Taken together, these results describe the interactions between the distribution of mutation effects, the competition-dependent selection and the energetic constraint and will help understand the emerging genetic and phenotypic diversity (see below).

### Number of coexisting genotypes

During the adaptation of an initially monomorphic population, *de novo* mutations generate polymorphism but at the same time purifying selection tends to reduce such diversity. How many genotypes are generated and maintained under competition for resources? Under the parameters explored in the simulations we observe that: after an initial burst of diversity, the mean number of genotypes first declines and later plateaus around a value below that obtained under neutrality (Fig. 3A). The drop in the mean number of genotypes is due to the energetic constraint, as populations evolving under neutrality or without such boundary do not suffer any decline (Fig. S4). When the populations’ phenotypes approach the energetic constraint, beneficial mutations become rarer and selection reduces the number of coexisting genotypes (Fig. S4). Despite the more abundant deleterious mutations, the populations can maintain a dynamic balance between the mutations that are purged by purifying selection and the newly emerging ones (inset of Fig. 3A).

**Figure 3.**
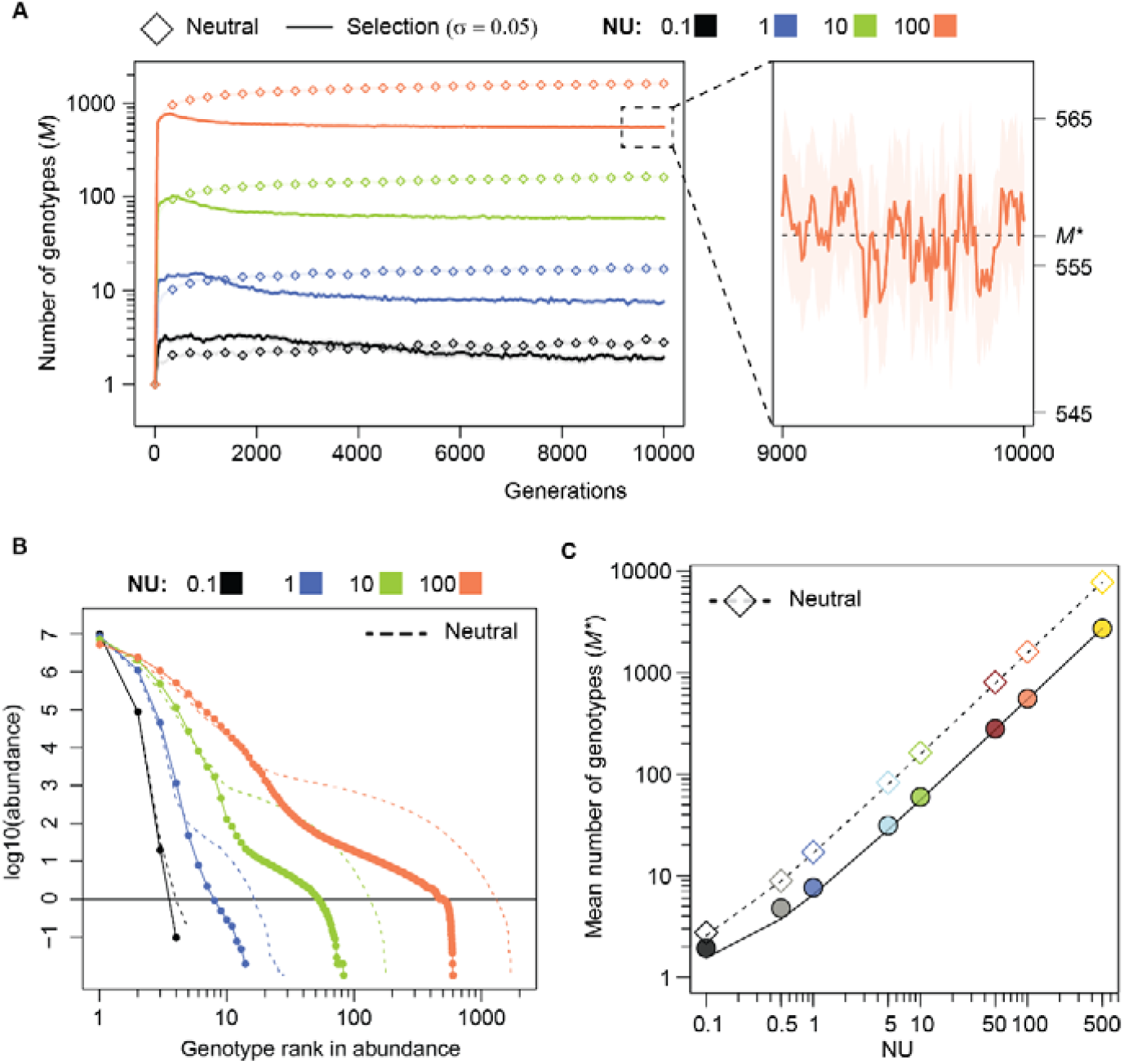
Genotypes’ dynamics and balance under competition for two resources. **A)** Number of genotypes present in the environment over time, under neutrality (diamonds) or under selection (lines). Lines represent the average across 100 populations and the shaded area their confidence interval. On the right, zoom over the last 1000 generation. Other parameters: *σ* = 0.05, *ρ* = 0 and *N* = 10^7^. **B)** Log-log scaled rank abundance distribution of genotypes at generation 10000. Dots and dashed lines represent the mean across 100 populations under selection or neutrality, respectively. Parameters as in A. **C)** Long-lasting number of genotypes, computed as the average over the last 1000 generations (e.g. dotted line in the right panel of A). The lines represent the linear regressions: *M*\* = *aNU* +1 where *NU*: (0.1,0.5,1,5,10,50,100,500} and *a* = (15.70 + 0.07,5.48 + 0.02} for the neutral or selection cases, respectively. Both axes are represented in *log* scale with ticks *every* (1,…, 9} · 10^*x*^. Other parameters as in A.

When we summarize the populations’ composition (at generation 10000) by calculating the average rank abundance distributions of the genotypes (instead of the species, Whittaker, 1965) we observe that few genotypes dominate the population (frequency above 1%) and the rest persists at low abundance (Fig. 3B). As the mutational input increases, the effect of selection becomes more pronounced as seen by stronger deviations from the rank abundance distribution observed under neutrality. As expected, under purifying selection less genotypes can be maintained at intermediate abundances (Haldane and Fisher, 1931; Wright, 1938). Under the conditions simulated here, populations maintain a dynamically stable genotype richness which increases linearly with the mutational input *NU,* and it’s about 1/3 of that expected under neutrality (with σ=0.05) (Fig. 3C). Thus, clonal populations with large *NU* and strong selection (*N σ* >>1) can maintain high levels of intraspecific variation under adaptation to few resources. However, we were unable to find an approximation to predict the number of genotypes, *M*\* across different combinations of *N, U* and *σ*, indeed *M*\*, deviates from *NU/σ* – the expected mean number of deleterious mutations under mutation-selection balance in a simple model of constant negative selection (Haigh, 1978).

### Population diversification into ecotypes

In this model, adapting populations consist of a cloud of many genotypes and we now characterize the phenotypic structures of these clouds. If a population consisted of a single genotype, the optimal strategy (hereafter Ω) would be to have the trait values that mirror the resource supply proportions 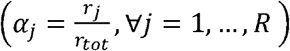, as represented by the star in Figs. 1–4. Such state cannot be invaded by any mutant 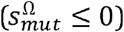, thus excluding diversity. However, if polymorphism already exists and metabolic trade-off is assumed, a large collection of types can stably coexist if they are distributed around Ω (Posfai, Taillefumier and Wingreen, 2017). Thus, we now ask the simple question: if there is no initial polymorphism and mutation is the only source of variation, will an initially maladapted population evolve a single strategy or multiple ones?

The simulations show alternative stable states: due to the stochastic nature of mutation, populations can evolve to either one or to multiple strategies, even if adapting under exactly the same conditions (e.g. Fig. 4A). Remarkably, the same ancestral genotype can give rise to many functionally similar genotypes (Fig. 4A, left panel), or can diversify into different ecotypes (Cohan, 2002): clusters of genotypes with distinct metabolic preferences, capable of coexisting indefinitely (Fig. 4A, right panel).

**Figure 4.**
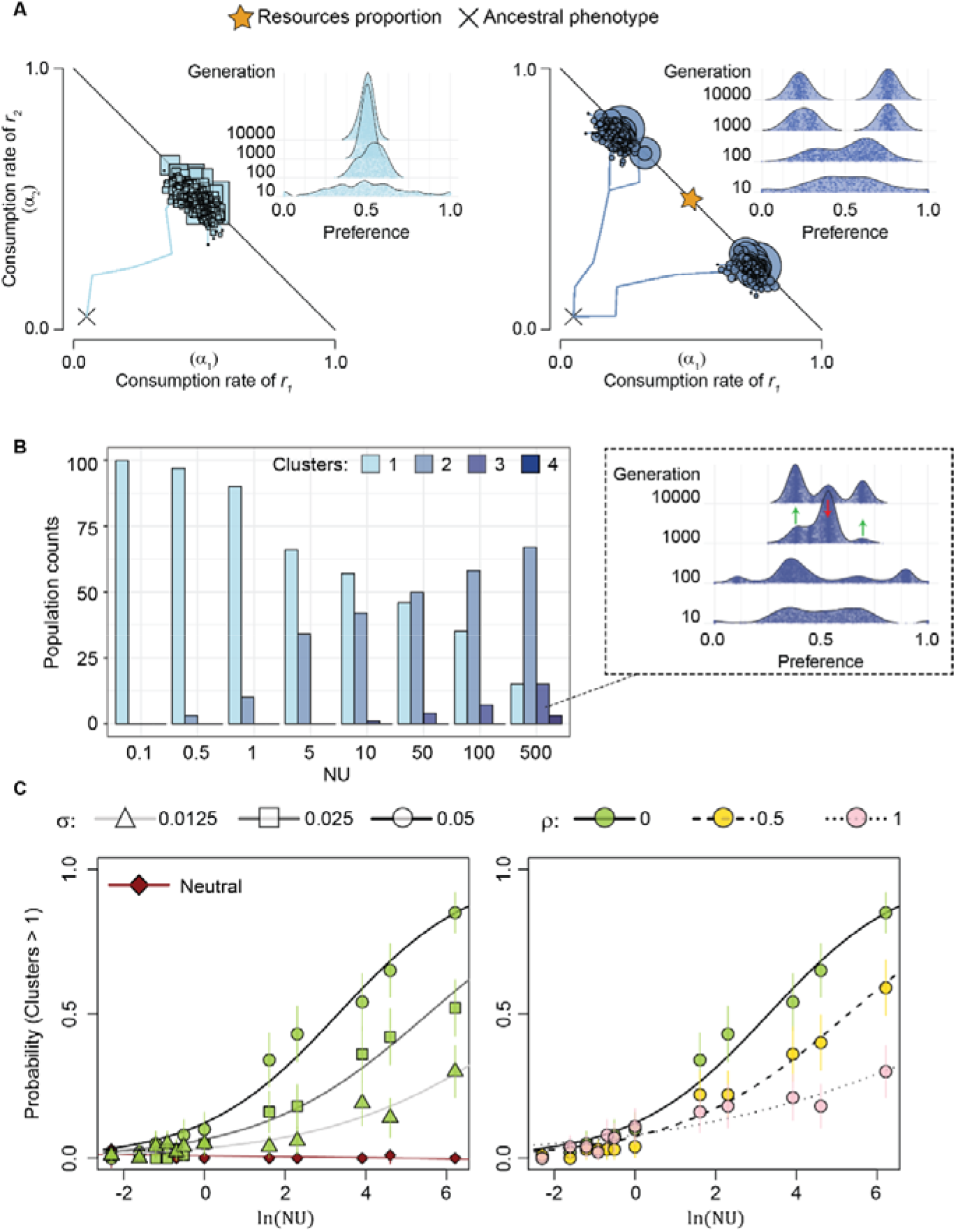
Ecological diversification under competition for two resources. **A)** Two example populations evolving under the same conditions (*N* = 10^7^, *U* = 10^-5^, *ρ* = 0.5, *σ* = 0.05, R = 2). The phenotypes and the preference distributions show one population that has evolved into a single optimal cluster (squares) and another population that gave rise to a stable diverse community composed by two clusters (circles). Lines connecting the shapes represent mutations. **B)** Counts of populations that evolved into 1,2 or more phenotypic clusters. Here, *N* = 10^7^, *σ* = 0.05, R = 2, ρ = 0. **C)** Populations diversify with a probability that increases with ln(*NU*) and σ but decreases with ρ. The lines represent the fit of the data to the logistic function: 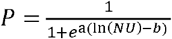, *NU*:{0.1,0.2,0.3,0.4,0.5,0.6,1,5,10,50,100,500}. The inferred parameters a and *b* are reported in Tables S1-2 and the full set of data is shown in Fig. S5. In the left plot: ρ = 0, while in the right one: σ = 0.05. The probabilities were computed as proportions out of 100 independent populations and their 95% confidence interval by normal approximation: 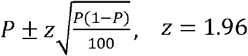.

We now investigate the conditions favoring such ecotype diversification, across several mutational inputs and in the regime of strong selection (*Nσ*□1). Using the mean shift clustering algorithm (Cheng, 1995) to group each adapted population into functional clusters (see details in *Methods* and Appendix S2), we find that the proportion of populations that evolved into *1,* 2 or more statistically distinct clusters, depends on the underlying mutation parameters. Under regimes of larger mutational input (i.e. more intense clonal interference) ecotypes with distinct preferences more likely emerge and coexist (Fig. 4B). Importantly, when *NU* is very large, the system can support supersaturation: the number of ecotypes inferred by the mean shift algorithm can exceed the number of limiting resources (Fig. 4B). This result is confirmed when using the k-means clustering algorithm (MacQueen, 1967) (data not shown). Although difficult to achieve, supersaturation is stable if the clusters’ fitness differs by a relatively small amount *ε* (with 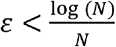) (Posfai, Taillefumier and Wingreen, 2017), but until then, the fate of supersaturated populations will be subjected to selection and *de novo* mutation (inset of Fig. 4B for an example of the fate of a specific population).

The probability of diversification (P) – computed as the proportion of populations that evolved more than one cluster – is close to zero when *NU*<1 but significantly increases when *NU≥1*. We summarize the increase observed in these simulations by the logistic function: 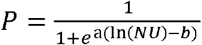 (see Fig. 4C and Tables S1-2). Note that the formation of ecotypes is not due to neutral processes as the populations adapting under neutrality did not diversify during the first 10000 generations simulated (Fig. 4C).

Not only the rate influences the diversification process (Fig. 4B) but also the type of mutations (Fig. 4C): under intense clonal interference, larger mutation effects (*σ*) and/or smaller pleiotropic effects (ρ) promote the formation of multiple ecotypes (Fig. 4C, Fig S5 and S6).

In summary, during the process of adaptation studied in these simulations, different qualitative outcomes can be observed: i) when the input of new mutations is low, the fittest genotype recurrently takes over as a cloud of genotypes until the population reaches a distribution around the generalist strategy that mirrors the resource supply (e.g. Video S1); ii) but when *NU* is large enough, the initial availability of many beneficial mutations causes adaptive radiation, opens the door for several genotypes to coevolve and for distinct ecotypes to stably coexist (e.g. Video S2).

### The genetic signature of diversification

We characterized the adapting populations by calculating their average pairwise genetic (*π_G_*) and phenotypic (*π_P_*) distances within populations. Both *π_G_* and *π_P_* increase with *NU* and always exceed the neutral simulations (Fig. 5A). However, genetic diversity does not necessarily imply functional diversity as some populations are observed to converge to similar phenotypes (see examples in Fig. S7) after an initial increase in diversity. Larger pleiotropy (*ρ*) in mutation effects reduces phenotypic diversity (Fig. 5A) and can foster a higher fraction of populations to exhibit phenotypic convergence. In the populations evolved with large *NU, π_P_* and *π_G_* correlate (*π_P_* ~0.7 *π_G_*, Fig. S8 top panel), but strong pleiotropy reduces the correlation (Fig. S8 bottom panel), explaining why fewer clusters are generated under this condition (Fig. 4C).

**Figure 5.**
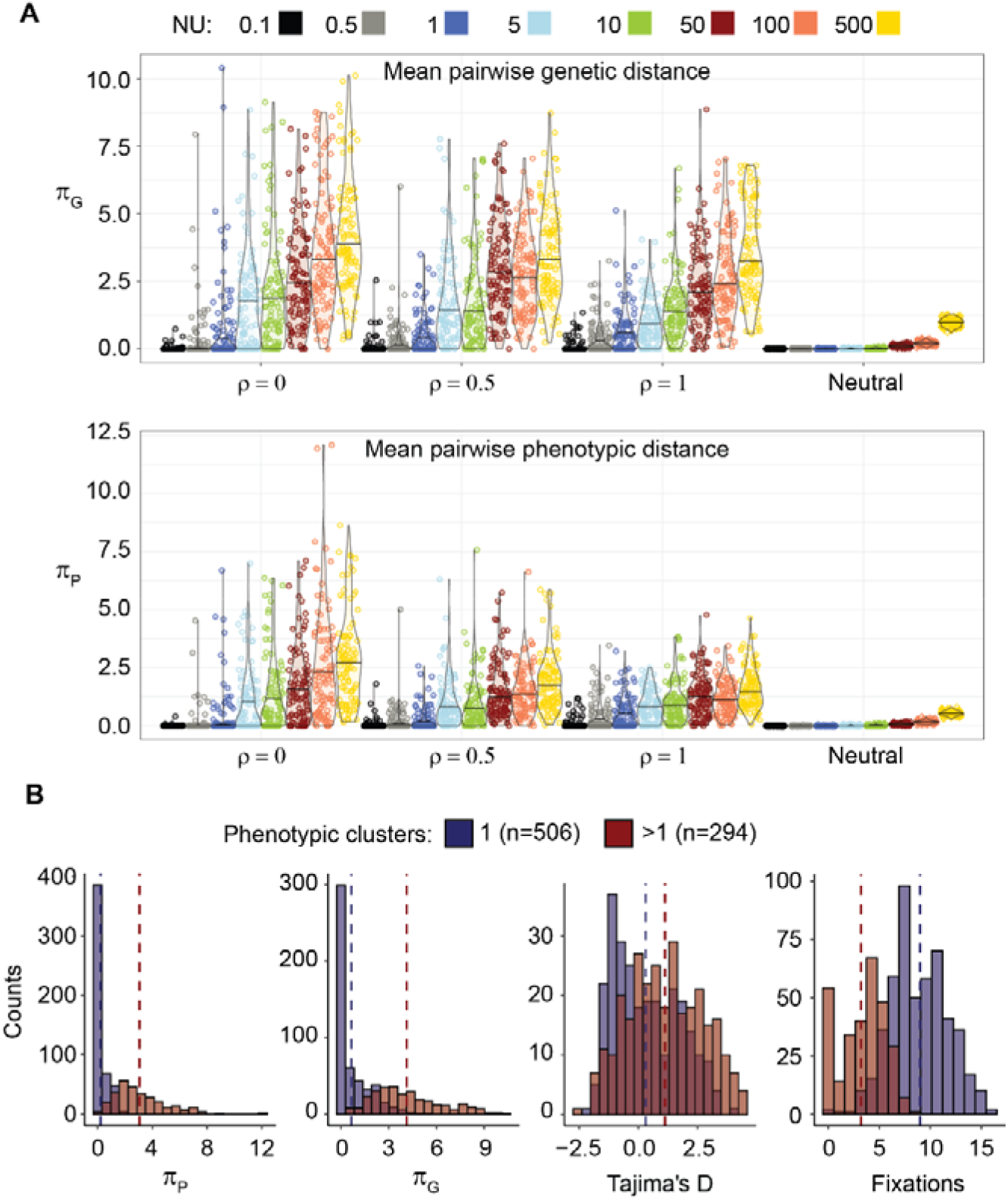
Phenotypic and genetic characterization of the evolved populations. **A)** Average pairwise genotypic (*π_G_*) and phenotypic (*π_P_*) diversity were measured within each population, as defined in the *Methods.* 100 independent populations were simulated for each of the conditions specified on the x axis and by the colors. Other parameters: σ = 0.05, 2 resources and *N* = 10^7^. **B)** π_P_, π_G_, Tajima’s D and fixations distributions of the populations that evolved into a single cluster (blue, *n*=506) or into multiple ones (red, n=294). Populations with different NU and ρ=0 were pulled together for a total of 800 populations. The dotted lines represent the means of the corresponding distribution.

Can we infer ecotype formation from the genetic composition of a population? The Tajima’s D statistic, that compares the pairwise genetic diversity with the number of segregating mutations in a sample, is meant to distinguish between different forms of selection. D is expected to be negative under recurrent sweeps or weak purifying selection, and positive under balancing selection (Tajima, 1989). We thus expect that the populations where stable ecotypes have formed show positive D values. Overall, the populations that diversified into multiple phenotypic clusters have on average larger *π_P_, π_G_*, Tajima’s D and fewer fixations (Fig. 5B, Mann-Whitney p-values < 10^-9^ for each of them). However, the distributions of Tajima’s D greatly overlap and require extra care in the study of out of equilibrium populations. In our simulations, under the accumulation of neutral mutations, larger mutational inputs push D towards negative values by generating larger numbers of segregating sites (Fig. S10). But, under selection, this can be counteracted by the increase of *π_G_*, and D presents a wider distribution and a non-monotonic relation with *NU* (Fig. S10). This makes the interpretation of the Tajima’s D more complex, and may help explain the overlap between the distributions of D in the simulation results of Fig. 5B.

The genetic diversity alone is better at discriminating the populations with multiple or single ecotypes (Fig. 5B), but its predictive power becomes less accurate in the regimes with larger pleiotropic effects (Fig. S9), due to the increased phenotypic convergence discussed above. Nevertheless, while with cross-sectional data it is difficult to establish that ecotypes have formed, with time series data the signal can be clearer (e.g. Fig. S7 where *π_G_* is large and D is positive along time). Statistics on population genomic data, together with functional measurements, should allow to identify populations that undergo ecological diversification.

### Small population size and accumulation of deleterious mutations can lead to speciation

So far, we have investigated adaptation in large populations with strong selection, where deleterious mutations hardly ever fix as most stay at low frequencies. Now, we explore simulations with different population sizes but under the same range of mutational inputs as before *(NU*=0. 1-500). When *N* = 10^5^, 10^7^ or 10^9^ the probability of diversification increases consistently (Table S3), confirming its relation with the parameter *NU* in regimes with large populations (Fig. 6A). Differently, when *N* = 10^3^ the probability of diversification shows a sharper increase (Fig. 6A, Table S3) as all the populations present multiple phenotypic clusters when *NU* ≥ 100. This is due to a different diversification process which requires the accumulation of deleterious mutations as we will explain. When *N* = 10^3^ and *NU* ≥ 100, we expect that deleterious mutations of a given effects can accumulate under the action of Muller’s ratchet (Felsenstein, 1974). In fact, in such regime, assuming constants = *σ* (here *σ* = 0.05), it follows that 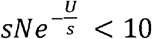, and the Muller’s ratchet should click in the time scale of our simulations (Gordo and Charlesworth, 2000). Contrarily, when *N* = 10^5^,10^7^ or 10^9^, it follows that 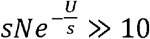, and the deleterious mutations should fix only in infinitesimally large time (Fig. S11). Furthermore, (Etheridge, Pfaffelhuber and Wakolbinger, 2012) showed that the ratchet can click when the parameter 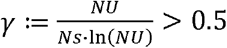, in a constants model. If we fix *s* = *σ* and *N* = 10^3^, this threshold condition occurs when ln(*NU*) > 4.8 (dotted line in Fig. 6A), consistent with the abrupt change in the observed probability of diversification and suggesting that when the ratchet turns, deleterious mutations will allow phenotypic clusters to form.

**Figure 6.**
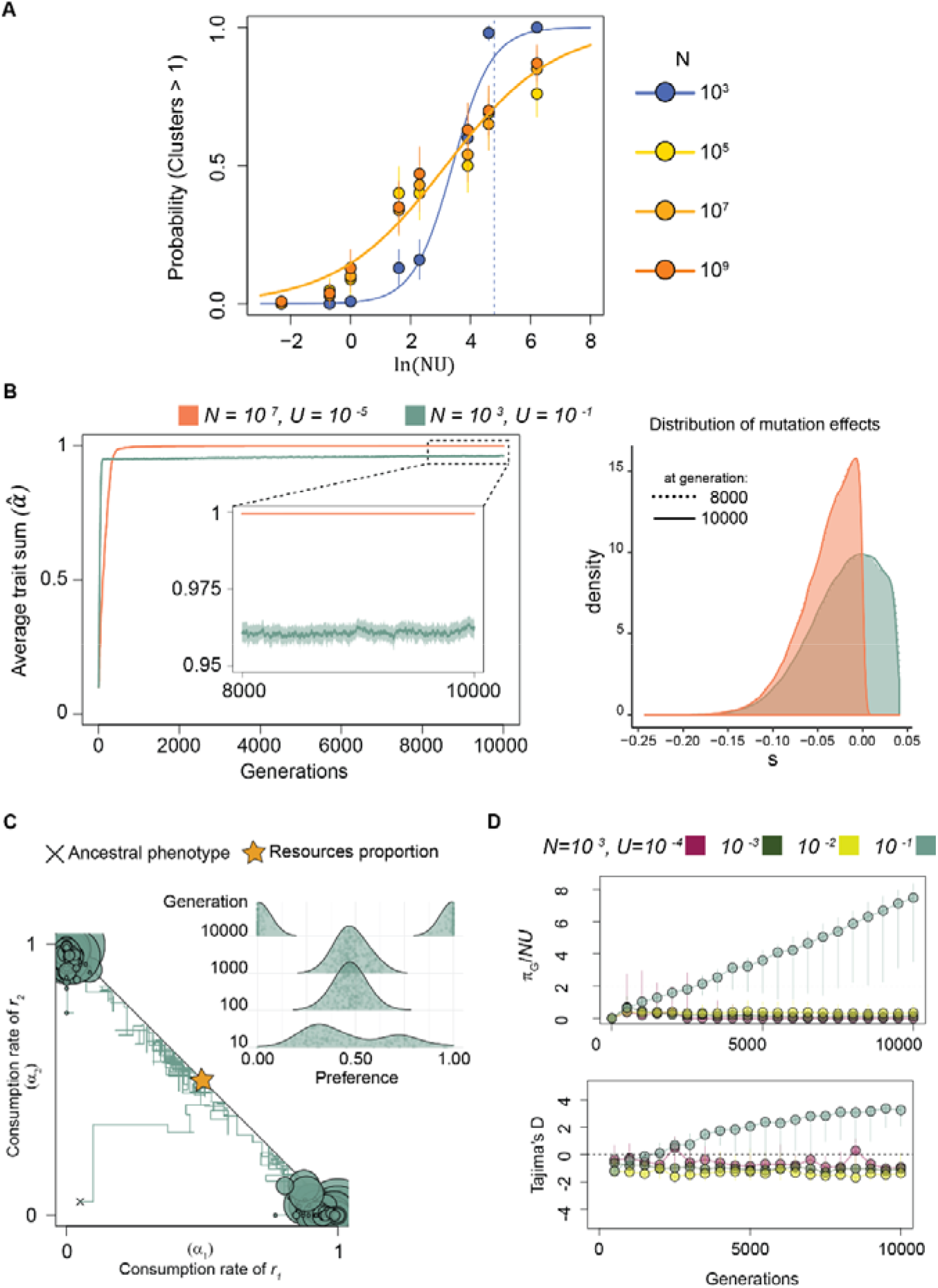
Speciation process in small populations with large mutational input. **A)** Probability of diversification across different population sizes. Continuous lines represent the fit of the data to the logistic function: 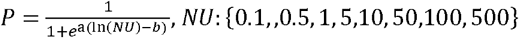, for *N* = 10^3^ [blue] or *N* = 10^5^, 10^7^, 10^9^ together [orange). The inferred parameters a and *b* are reported in Tables S3. The dotted line represents the threshold (ln(*NU*) = 4.78) when 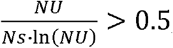, *N* = 10^3^ and 10 *s* = *σ*. The probabilities were computed as for Fig. 4. **B)** Average trait sum over time and distribution of mutation effects at generations 8000 or 10000. **C)** Example population adapted with *N* = 10^3^, *U* = 10^-1^, *ρ* = 0, *σ* = 0.05, *R* = 2 for 10000 generations. The inset shows the preference distribution at generations 10, 100, 1000 and 10000. **D)** *ρ_G_*/*NU* and Tajima’s D over time. Circles represent the median, while vertical bars range from the 25^th^ to the 75^th^ quartiles. The dotted lines represent the expected value at neutral equilibrium. In particular *π_G_* = 2*NU* and Tajima’s D=0. Other parameters: σ = 0.05, ρ = 0.

Studying large and small populations with different mutation rates (*U*=10^-5^ or *U*=10^-1^, respectively) but equal mutational input (*NU*=100), we compare the two qualitatively different regimes. Fig. 6B shows how the ratchet reduces the average trait sum of the small populations relative to the large populations (case where the ratchet does not turn). Because in this parameter range beneficial mutations are still common (Fig 6B, right panel), their effect balances that of the deleterious mutations and the mean fitness equilibrates at intermediate levels (as in Goyal *et al*., 2012). While the mean fitness is constant, the phenotypic distribution becomes bimodal. Contrarily to what was previously observed in the large populations (diversification in the early steps of adaptation and then long-term stabilization), in these smaller populations *(N* = 10^3^) with large mutation rate (*U* = 10^-1^), diversification happens after the energetic boundary in reached (Fig. 6C). The continuous accumulation of deleterious and compensatory mutations drives the clusters apart until they reach the specialist extremes and finally stabilize (see an example in Fig. 6C and Fig. S12). Thus, in the small populations with large mutational inputs, ecological diversification can maximize the functional diversity within the population, lead to a continuous increase of genetic diversity and push the Tajima’s D values well above neutral expectations (Fig. 6D). This process should in principle lead to incipient speciation in the long run.

## Discussion

Microbial communities are vital for humans and many other host species (Nicholson *et al*., 2012; Sunagawa *et al.,* 2015). Emerging observations of evolution in such ecosystems (Barroso-Batista *et al*., 2014; Garud *et al*., 2019; Zhao *et al*., 2019) motivate new theories where the mechanisms that generate diversity involve complex forms of selection and clonal interference (Gordo, 2019). We propose that a simple eco-evolutionary model of resource competition, describing the mechanisms behind ecological divergence, can help understand diversity within ecosystems. This framework can be generalized to incorporate other evolutionary mechanisms, such as other forms of selection (Good, Martis and Hallatschek, 2018), transmission and horizontal gene transfer, and can serve to bridge an existing gap between ecology and population genetics.

Population genetics models of clonal interference have greatly advanced our understanding of microbial adaptation (Gerrish and Lenski, 1998; Park and Krug, 2007; Good *et al*., 2012; de Sousa *et al*., 2016). However, clonal interference is rarely considered in theoretical studies of ecosystems (Farahpour *et al*., 2018), even though it greatly impacts the evolution of microbes within real communities (Barroso-Batista *et al*., 2014) and may be relevant in key ecosystems such as the human microbiota (Zhao *et al*., 2019). Commensal species in the gut have large population sizes ~10^8^ cells/g. If each bacterium mutates in the gut as it does in the laboratory (Drake, 1991), then each gram of material will host around 10^5^ new mutant cells every generation. Even if only 0.1% brings up a benefit (Perfeito *et al*., 2007), clonal interference still extensively affects the gut microbiota dynamics.

Here we have studied an ecological model where clones do not compete for fixation but for resources. Modeling competition explicitly allows to make testable predictions about different measures of diversity as both the traits and genomes can now easily be studied. We show that clonal interactions can drive an initial monomorphic population to polymorphism with distinct ecotypes, deviating from the simple expectation of adapting to a single optimal phenotype (Fig. 4 and Fig. 6).

Taking the MacArthur model, it was previously demonstrated that metabolic trade-offs promote coexistence of more species than resource types, thus overcoming the competitive exclusion principle (Posfai, Taillefumier and Wingreen, 2017). Extensions of this framework already demonstrated its power in recapitulating experimental results from studies of soil, plant (Goldford *et al*., 2018) or even mammalian gut microbiotas (Leónidas Cardoso *et al*., 2020). And further extensions provided new analytical results of how populations can adapt under competition for resources and demonstrated that directional selection can limit ecological diversification (Good, Martis and Hallatschek, 2018). This pattern is also observed in our simulations as we find that stronger pleiotropy, which causes stronger directional selection, limits the emergence of clusters (Fig. 4C). Our work differs from the latter by focusing on mutations that alter the metabolic strategies and by quantifying the phenomena that emerge out of equilibrium. We show that under clonal interference, the outcome of phenotypic adaptation is probabilistic, whereby the populations can evolve paths that were not observed under equilibrium assumptions. We further observe that small populations with large mutation rates will diversify by the accumulation of deleterious mutations and generate different species-like lineages.

Trade-offs are commonly assumed and expected to affect evolutionary trajectories (Farahpour *et al*., 2018; Amado and Campos, 2019), but this is not always observed. While many empirical results have confirmed the role of trade-offs during adaptation (Bell and Reboud, 1997; Bull, Badgett and Wichman, 2000; Turner and Elena, 2000; Dykhuizen and Dean, 2004; Greene *et al*., 2005; Duffy, Turner and Burch, 2006; Coffey *et al*., 2008; Ward, Perron and MacLean, 2009; Bailey and Kassen, 2012; Li, Petrov and Sherlock, 2019), others did not find evidence for any (Reboud and Bell, 1997; Kassen and Bell, 1998; Turner and Elena, 2000; Trindade *et al*., 2009; Bedhomme, Lafforgue and Elena, 2012). Here we assumed a linear trade-off in the form of an energetic constraint, which only affects well adapted genotypes. Thus, in our model the observation of a trade-off depends on the time at which it is measured. It would be interesting to test for trade-offs at different times during adaptation, as this could explain some of the contrasting findings outlined above. Compatible with this hypothesis, a trade-off in *Escherichia coli* ability to grow in the presence of both glucose and lactose was found, but it only emerged after a period of constraint-free adaptation (Satterwhite and Cooper, 2015). We find that in large populations, the metabolic tradeoff in the resource consumption is not required for the formation of distinct ecotypes but it promotes their stable coexistence (Fig. 4, S7 and Video S2).

In many ecosystems, the coexisting types seem to outnumber the limiting resources, and, solving this contention has motivated numerous studies. Previous eco-evolutionary analysis suggest that adding evolutionary changes confirms (Edwards *et al*., 2018) or even exacerbates (Shoresh, Hegreness and Kishony, 2008) this paradox. Perhaps surprisingly, our simulations show that large mutational inputs maintain a dynamically stable number of types that overcome the competitive exclusion (Fig. 3). And at the functional level, diversity generally respects the exclusion principle (number of types ≤ number of resources) but with exceptions: in a regime of strong clonal interference, the number of extant ecotypes can be larger than the number of limiting resources (Fig. 4B).

We show that the process leading to ecological diversification is strongly influenced by the underlying molecular parameters: regimes of low clonal interference *(NU* <1) lead to the evolution of a single generalist population, but large mutational inputs *(NU≥1)* more likely lead to the formation of two or more differentially specialized ecotypes (Fig. 4 and Video S1-S2). Our analysis demonstrates that large mutation effects and weak pleiotropy foster ecological diversification (Fig. 4C). Different pleiotropic effects are meant to represent different interactions between the traits under selection. If the available resources are similar (e.g. chemical composition) and/or the metabolic processes involved in their consumption share many genes, this could increase the chances that a mutation affects the two traits simultaneously leading to large pleiotropy. Contrarily, less related resources could involve more independent effects leading to smaller pleiotropy, promoting diversification (Fig. 4C). This interpretation could explain why adaptive diversification occurred in some experimental evolution setups (Friesen *et al*., 2004; Sandberg *et al.* 2017) but was not observed in others (Satterwhite and Cooper, 2015; Sandberg *et al.* 2017). In agreement with this hypothesis, Sandberg and colleagues showed that evolving on less metabolically related resources promoted ecological diversification (Sandberg *et al.* 2017).

Sympatric diversification can be observed experimentally and predicted by theoretical models (Friesen *et al*., 2004). The framework of adaptive dynamics has been extensively used in this context as it describes evolution on fitness landscapes that change dynamically due to frequency-dependent interactions (Geritz *et al*., 1998; Doebeli, 2011). However, these models lack the explicit mechanism of competition for resources and are based on equilibrium assumptions: the populations first evolve to an equilibrium state before diversification occurs, as explained by the concept of evolutionary branching points. Our individual-based model, in contrast, allows to follow adaptation of populations that undergo strong non-equilibrium dynamics and can accumulate deleterious mutations if they are small (Fig. 6). Previous studies (Rosindell, Harmon and Etienne, 2015; Ispolatov, Madhok and Doebeli, 2016) highlighted how considering evolution at the individual level is necessary to fully understand the adaptation process.

We find that the genetic diversity of a population can be used to predict the underlying phenotypic structure, but with limitations (Fig. 5). Other statistics based on genetic data (such as the Tajima’s D) can help in understanding the diversity structure, especially if these are assayed along time (Fig. 6).

The model studied here serves as a first step at integrating phenotypic and genetic data in a relatively simple environment It predicts that the typical high mutational input of bacterial species and cancer cells, coupled with an energetic constraint, is a mechanism capable of generating functionally diverse clonal communities. Future frameworks addressing microbial ecology and evolution will need to address how space, migration and/or fluctuating conditions affect the patterns of diversity. In addition, cooperation between genotypes (such as cross-feeding or use of costly public goods) and many more resources have the potential to shape diversity levels and should also be investigated in future studies.

## Author contributions

MA and IG designed the study, all authors wrote the manuscript and provided final approval for publication.

## Acknowledgements

The authors acknowledge C Bank and the members of the Evolutionary Dynamics and Evolutionary Biology labs of the IGC for their assistance throughout the development of this work; PR Campos, T Paixão, S Miller, G Sgarlata and R Ramiro for their comments on the manuscript This work was supported by Portuguese Science Foundation (FCT) Grant PTDC/BIA-EVL/31528/2017; Deutsche Forschungs-gemeinschaft (DFG) Grant SFB 1310; and a cooperation agreement between University of Cologne and Gulbenk?an Institute. MA was supported by FCT Grant PD/BD/138735/2018.

Data archival location will be made public upon acceptance.

