## Supplementary Information for "Molecular signatures of resource competition: clonal interference favors ecological diversification and can lead to incipient speciation"

**Supporting Information**

**Appendix S1 – The strength of selection under competition for substitutable resources**

We model adaptation under competition for substitutable resources. The ecological dynamics follow (Posfai, Taillefumier and Wingreen, 2017) and dictate how selection acts on emerging mutants. To understand how such competition-dependent selection changes along the adaptive process we first try an analytical approach. Considering that a population at time $t$ contains $M$ types which compete for $R$ resources, then selection acting on each type $i$ is given by:

$s_{i}(t)=\sum_{j=1}^{R} \frac{\alpha_{j}^{(i)}r_{j}}{\sum_{k=1}^{M(t)} n_{k}(t)\cdot\alpha_{j}^{(k)}}-\delta$	(1)

where $\alpha_{j}^{(k)}$ represents the consumption rate of resource $j$ by type $k$, $n_{k}$ is its density, $r_{j}$ is the concentration of resource $j$ and $\delta$ is the death rate. The denominator of (1), which we will refer to as $e_{j}^{(M)}:=\sum_{k=1}^{M} n_{k}\cdot\alpha_{j}^{\left( k \right)}$, represents the population investment on each resource $j$. Let us further simplify the problem by considering a monomorphic ($M=1$) population whose phenotypes ($\vec{\alpha}$) mirror the resource input proportions:

$\frac{\alpha_{j}}{\sum_{k}^{R} \alpha_{k}}=\frac{r_{j}}{\sum_{k}^{R} r_{k}},\forall j=1,\ldots,R$.

This strategy consists in the optimal one for a given energetic investment ($\sum_{k}^{R} \alpha_{k}$).

It follows that the selection acting on an emerging mutant, $mut$, is:

$s_{mut}(\vec{\alpha},\vec{\Delta})=\frac{\sum_{j}^{R} \Delta_{j}}{\sum_{j}^{R} \alpha_{j}}$ (see main text).

This represents the so-called invasion selection that acts on the low abundant mutants ($n_{mut}\ll N$). In order to retrieve an explicit form of selection, we made use of the continuous univariate distribution theory (Johnson *et al.*, 1994). We know that, by assumption, the phenotypic changes $\Delta_{j}$ are normally distributed ($N(0,\sigma)$) and so are their additive effect: $\sum_{j}^{R} \Delta_{j}\sim N(0,\sigma^{*})$ where $\sigma^{*}=\sigma\sqrt{(R-1)\cdot\rho^{2}+1}$ (see *Methods* in the main text). However, due to our energy constraint assumption $(\sum_{j=1}^{R} \alpha_{j}^{(i)}\leq1$ , $\forall i=1,\ldots,M)$, the mutations are conditional to $-\infty<\sum_{j}^{R} \Delta_{j}\leq1-\sum_{j}^{R} \alpha_{j}$ and the distribution of beneficial mutations is, in fact, a two-sided truncated normal distribution:

$$\sum_{j}^{R} \Delta_{j}\sim N\left( 0,\sigma^{*} \right), 0<\sum_{j}^{R} \Delta_{j}\leq1-\sum_{j}^{R} \alpha_{j}$$

From (Johnson *et al.*, 1994), if $X\sim N\left( \mu,\sigma\right)$ then:

$$E\left[ X| a<X<b \right]=\mu+\sigma\frac{\varphi\left( A \right)-\varphi\left( B \right)}{\Phi\left( B \right)- \Phi\left( A \right)}$$

With $A:=(a-\mu)/\sigma$, $B:=(b-\mu)/\sigma$, $\varphi\left( x \right):=\frac{1}{\sqrt{2\pi}}e^{-\frac{1}{2}x^{2}}$ and $\Phi\left( x \right):=\frac{1}{2}\left( 1+erf(\frac{x}{\sqrt{2}}) \right)$, with $\mathrm{erf} \left( x \right):=\frac{1}{\sqrt{\pi}}\int_{-x}^{x} e^{-t^{2}}dt$. Consequently, in our case:

$$E\left[ \sum_{j}^{R} \Delta_{j}| 0<\sum_{j}^{R} \Delta_{j}\leq1-\sum_{j}^{R} \alpha_{j} \right]=\sigma^{*}\left( \frac{\frac{1}{\sqrt{2\pi}}-\varphi\left( \frac{1-\sum_{j}^{R} \alpha_{j}}{\sigma^{*}} \right)}{\Phi\left( \frac{1-\sum_{j}^{R} \alpha_{j}}{\sigma^{*}} \right)-\frac{1}{2}} \right)$$

From this we computed the corresponding expected selection coefficients as

$E\left[ s^{+} \right]=\frac{E\left[ \sum_{j}^{R} \Delta_{j} \right]}{\sum_{j}^{R} \alpha_{j}}$	(2)

over different values of $\sum_{j}^{R} \alpha_{j}$ (see Fig. 2A in the main text).

Knowing how selection varies over the adaptation process allows to predict what the expected effect of beneficial mutations is (see Fig. 2B in the main text) but also provides the relation between selection and time. In fact, the integral of (2) serves as scale of the adaptation rates across different conditions (see Fig. 2C).

**Appendix S2 – Ecotype call via the *mean shift* clustering algorithm**

At the end of the simulated adaptations (after 10000 generations), each independent population $x$ consists of a collection of $M_{x}$ types, each of them defined by a phenotypic vector $(\alpha_{1}^{\left( i_{x} \right)},\alpha_{2}^{(i_{x})})$ with $i_{x}=1,\ldots, M_{x}$. For each of the evolved populations, in order to identify functional clusters of genotypes, we tested whether the phenotypes followed a unimodal or a multimodal distribution via the mean shift clustering algorithm (Cheng, 1995), implemented through the *R* package *meanShiftR* version 0.53 (Lisic, 2018). The latter performs classification of the set of phenotypes using steepest ascent to local maxima in a kernel density estimate. We used a two-dimensional Gaussian kernel (default) with standard deviation (“*bandwidth”* parameter) equal to $\sigma$ on each dimension and we set $\sigma^{*}$ to be the minimum distance between distinct clusters (“*epsilonCluster”* parameter) as the effective mutational step $\sigma^{*}$ increases with $\rho$. The remaining parameters were as default settings. For each population we collect the number of detected clusters.

**Supplementary figures**

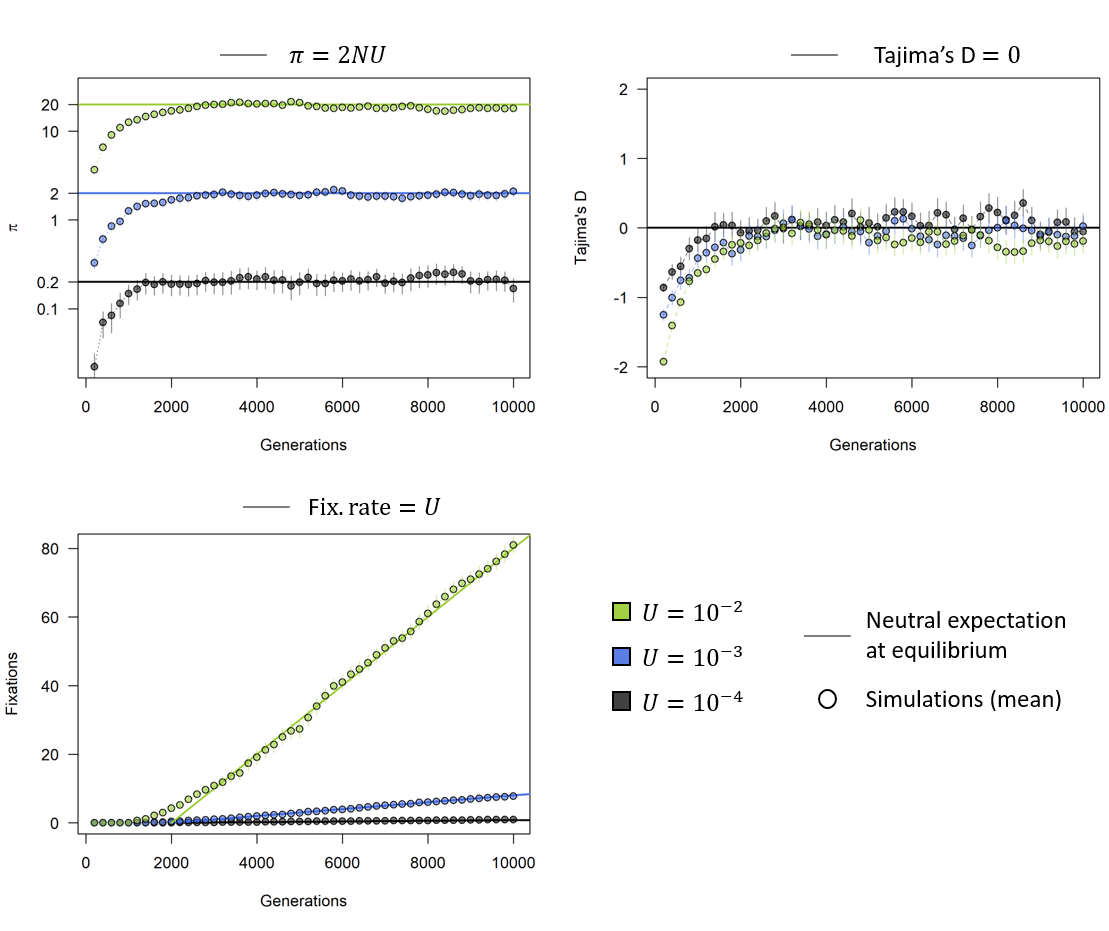

**Figure S1. Code validation.** In order to test the validity of the code, we can compare simulation outcomes with well-known analytical expectations of genetic quantities. Under neutrality, the expected pairwise genetic difference within populations ($\pi_{G}$) is *2NU*, the expected Tajima’s D is 0 and mutations should fix at rate *U*. Lines resent these predictions and circles represent the observations (mean and standard error). Here$N={10}^{3}.$

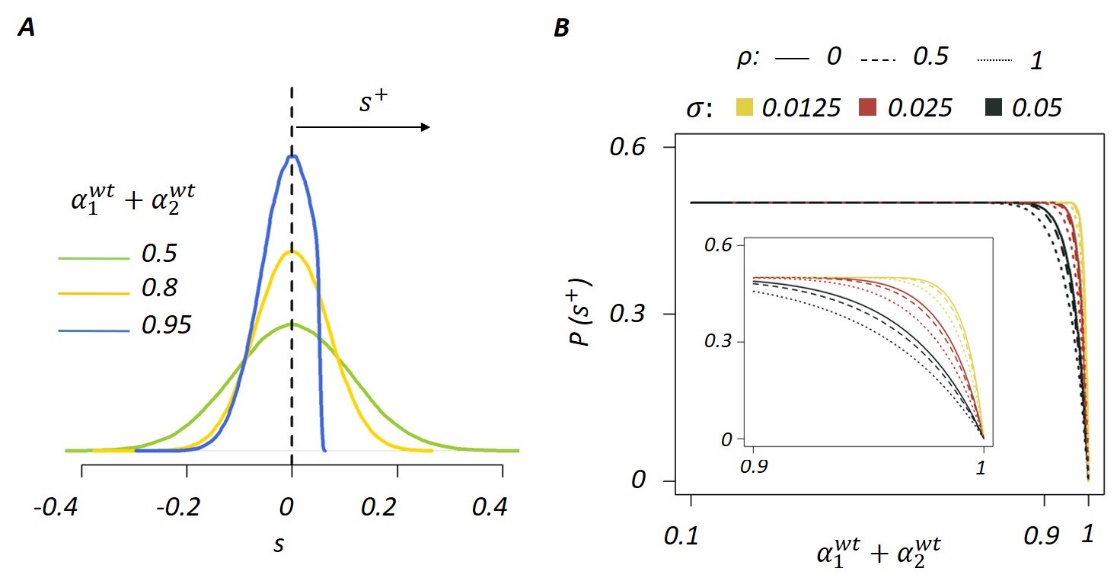

**Figure S2. Beneficial mutations become rare in well adapted populations. A)** Distributions of selection coefficients in differentially adapted monomorphic populations. Three examples of adapted populations are taken with traits $, \sum\alpha$= {0.5,0.8,0.95}, σ =0.05 and ρ = 0.5. **B)** Analytical approximation of the proportion of beneficial mutations. The probability of experiencing a beneficial mutation, computed as $P(s^{+})=\frac{\int_{0}^{(1-(\alpha_{1}^{wt}+\alpha_{2}^{wt}))} N(0,\sigma*)}{\int_{-\infty}^{(1-(\alpha_{1}^{wt}+\alpha_{2}^{wt}))} N(0,\sigma*)}$, is plotted against the trait sum of the parental wild-type, $\alpha_{1}^{wt}+\alpha_{2}^{wt}$. The inset shows the effect of the energetic constraint.

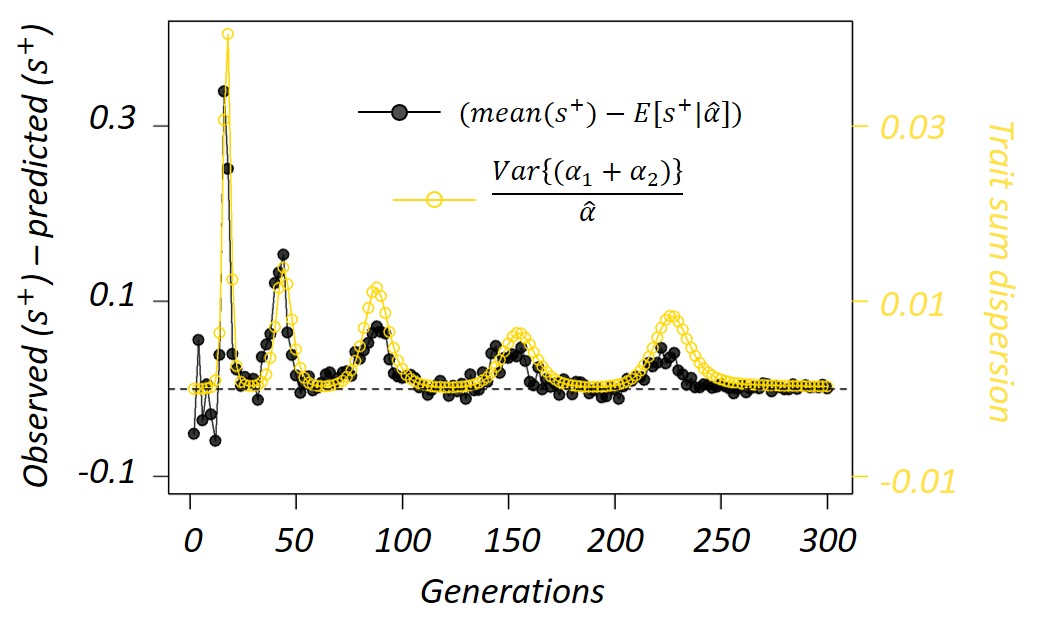

**Figure S3. Deviations from the analytical expectation are predicted by the population trait dispersion.** We compare the deviation from the predicted mean positive selection (shown in black) and the dispersion index of the population trait sums, computed as $\frac{Var\{(\alpha_{1}+\alpha_{2})\}}{\hat{\alpha}(t)}=\frac{1}{\hat{\alpha}(t)}\cdot\frac{\sum_{i}^{M(t)} n_{i}(t)\left( {(\alpha}_{1}^{(i)}+\alpha_{2}^{(i)})-\hat{\alpha}(t) \right)^{2}}{N-1}$. Peaks of stronger selection co-occur with dispersion peaks. The data shown here refer only to the black population from Fig. 2B but the same pattern is found in all the other populations.

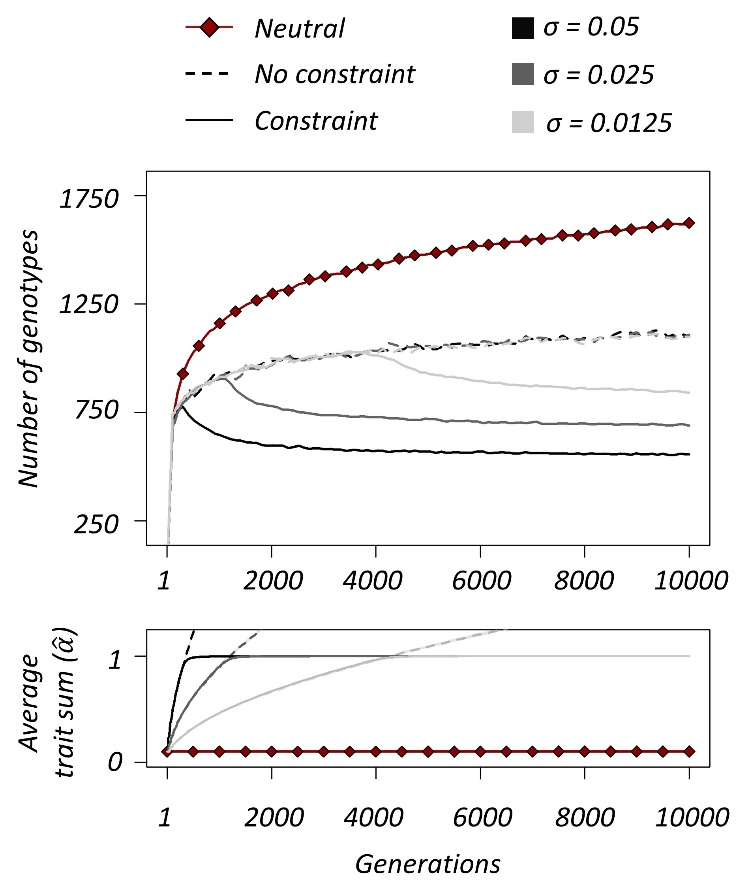

**Figure S4. Mean number of genotypes and the dependence on the energetic constraint**. Average number of genotypes and mean population phenotype over time, under different regimes. The continuous lines are populations evolving in the framework described in the main text (selection + energy constraint). The dotted lines and the red diamonds, show the populations evolving under selection but with no constraint and in the absence of selection, respectively. When the mean population phenotype approaches the energy constraint $(\hat{\alpha}=1)$ (bottom panel), the number of genotypes decreases (top panel). Interestingly, before (or in the absence of) the energy constraint, the number of genotypes does not depend on the mutation effect σ. Other parameters: $R=2, N={10}^{7}, U={10}^{-5}, \rho=0.$ Lines are average over 100 simulations.

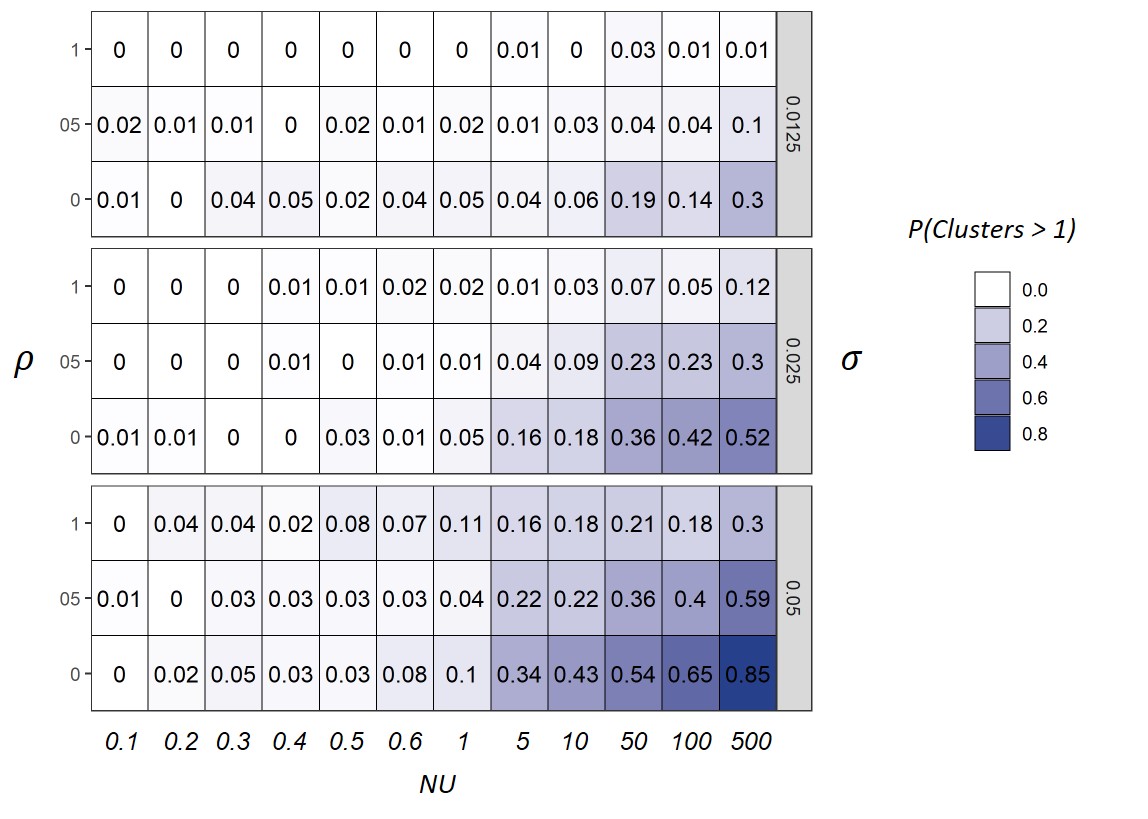

**Figure S5**. **Diversification probability across different parameters combinations.** 100 populations adapted under 108 parameter combinations. Here we report the probability of phenotypic diversification (as defined in the main text) for each of them. Darker blue indicates higher probability, which can be obtained via increasing NU and/or σ or decreasing ρ.

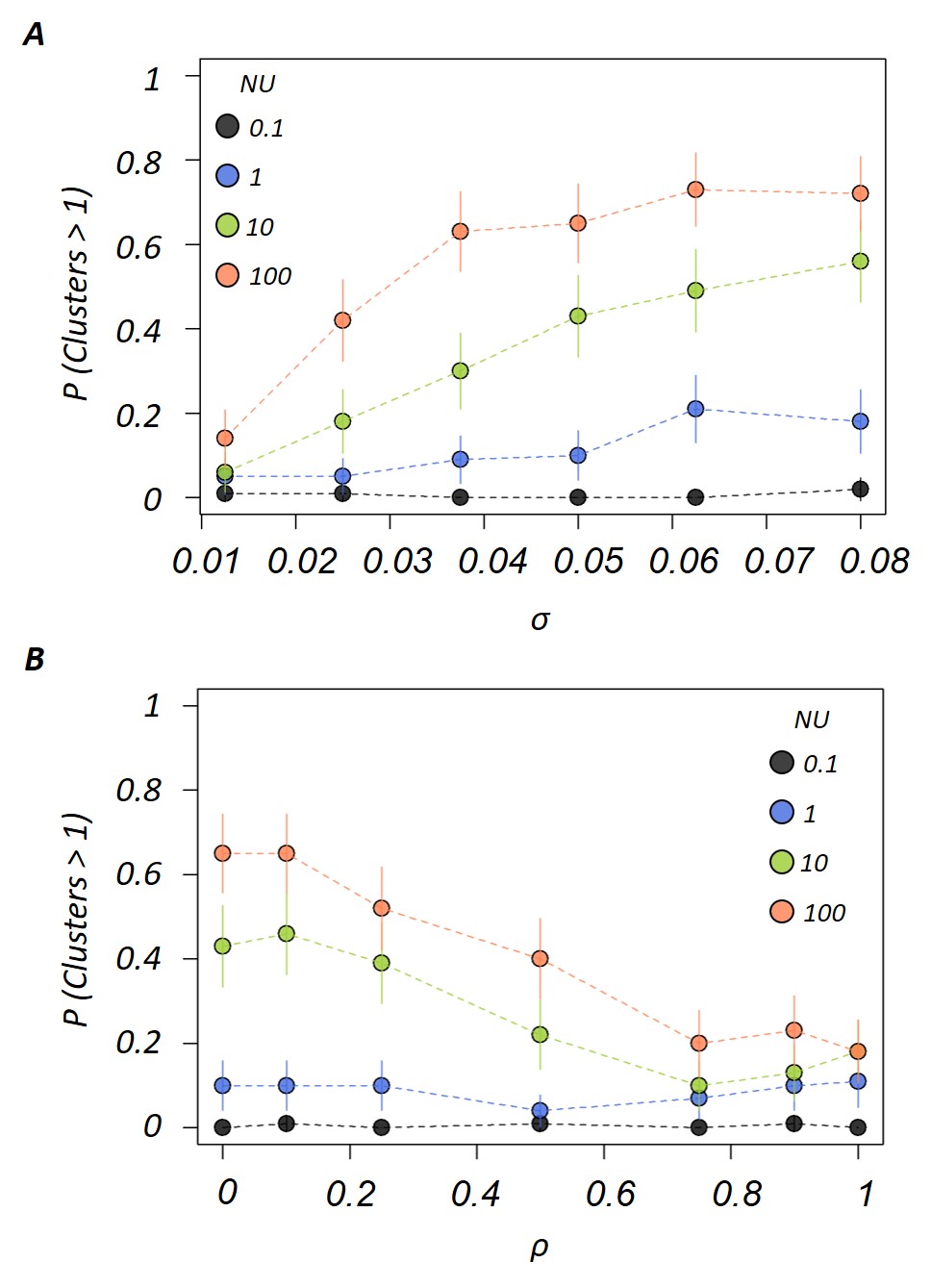

**Figure S6. The probability of diversification depends on** $\boldsymbol{\sigma}$ **and ρ.** Each dot represents the proportion of populations that evolved into more than one cluster. **A)** Under intense clonal interference, $P$ increases with $\sigma$ until a maximum at intermediate $\sigma$. Other parameters: $\sigma:\left\{ 0.0125,0.025,0.0375, 0.05,0.0625, 0.8 \right\}, \rho=0$. **B)** Under intense clonal interference, $P$ is larger for smaller $\rho$ and vice versa. Other parameters: $\rho:\left\{ 0,0.1,0.25, 0.5,0.75,0.9, 1 \right\},\sigma=0.05$.

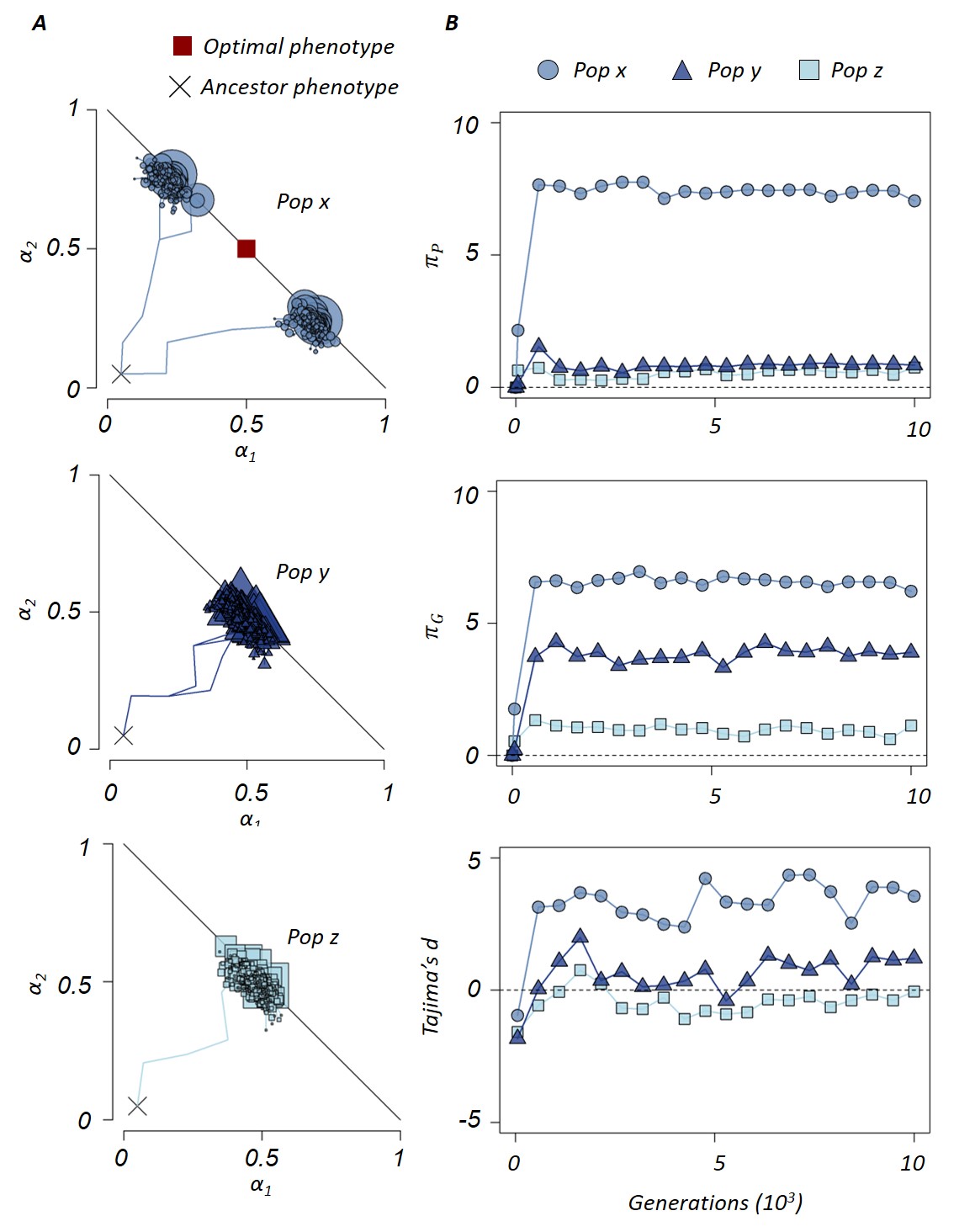

**Figure S7. Three different outcomes of adaptation.** Three example populations which evolved under equal conditions but resulted in contrasting outcomes: population x (circles) evolved two phenotypically and genotypically distinct ecotypes, population y (triangles) evolved functionally convergent lineages, while population z (squares) evolved as a single functional and genetic lineage around the optimum. **A)** Adaptation walks on the phenotypic space of the three populations. Each shape represents the mapping of a genotype on the phenotypic space and its size is proportional to the abundance in the population. The lines connecting the dots represent the mutations and are shown only for the genotypes that are still present (shapes). **B)** Average phenotypic and genetic distances and Tajima’s D over time of the three populations. Other parameters $\sigma= 0.05, \rho=0.5, N={10}^{7}$and U$={10}^{-5}$.

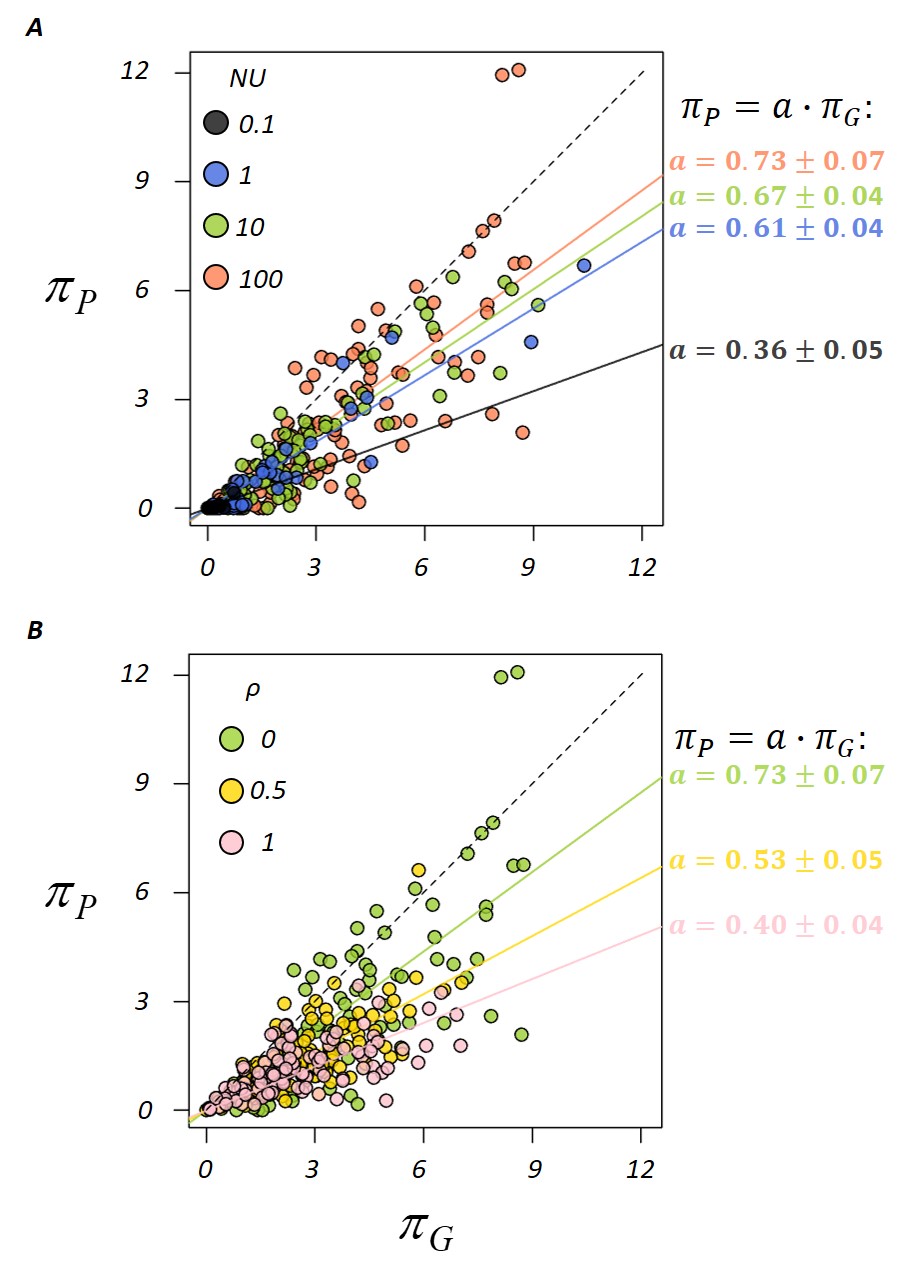

**Figure S8. Genotype to phenotype diversity.** Correlation between the phenotypic ${(\pi}_{P})$ and the genotypic ${(\pi}_{G})$ diversity within populations. Each dot represents the $\pi_{P}$ and $\pi_{G}$ of a population sampled at the end of the simulation (generation 10000). The continuous lines represent the linear regressions ${(\pi}_{P}{=a\cdot\pi}_{G})$ whose slopes, $a$, are reported on the figure. **A)** Phenotypic diversification increases with $\mathrm{NU}$. Other parameters: $\sigma= 0.05, \rho=0.$ **B)** Phenotypic diversification increases with $\rho$. Other parameters: $\sigma= 0.05, NU=100.$

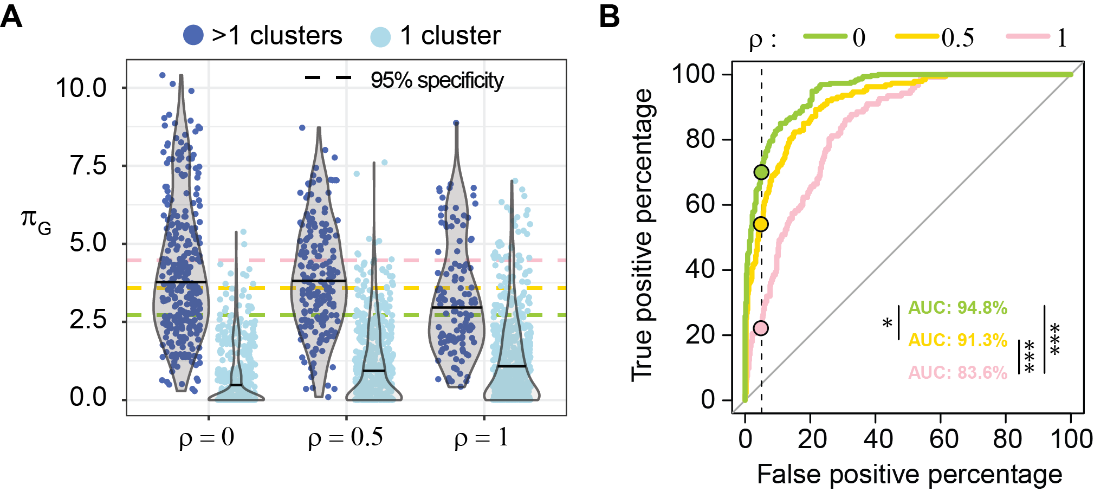

**Figure S9. Ecotype prediction by the genetic diversity. A)** $\pi_{G}$ distributions of the populations that evolved in more than one cluster (dark blue) or in a single cluster (light blue). Data form Fig. 5A with different NU were pulled together for a total of 800 populations per condition (ρ). The violin plots show the distribution of the data and their median. The dotted lines represent the $\pi_{G}$ threshold that would ensure 95% specificity in ecotype prediction (i.e. 5% false positive) and correspond to the circles in panel B. **B)** $\pi_{G}$ was used to predict whether each population is composed by one or multiple ecotypes. The ROC curves represent the sensitivity over one minus specificity of the prediction outcomes, for varying thresholds. The dotted line and the circles represent the thresholds that ensure 5% false positive rate and these are: $\pi_{G}>2.7, 3.6$or $4.5$ for ρ=0, 0.5 or 1, respectively. The area under the curve (AUC) is reported on the figure. * or *** indicate p-value <0.01 or <0.0001, respectively.

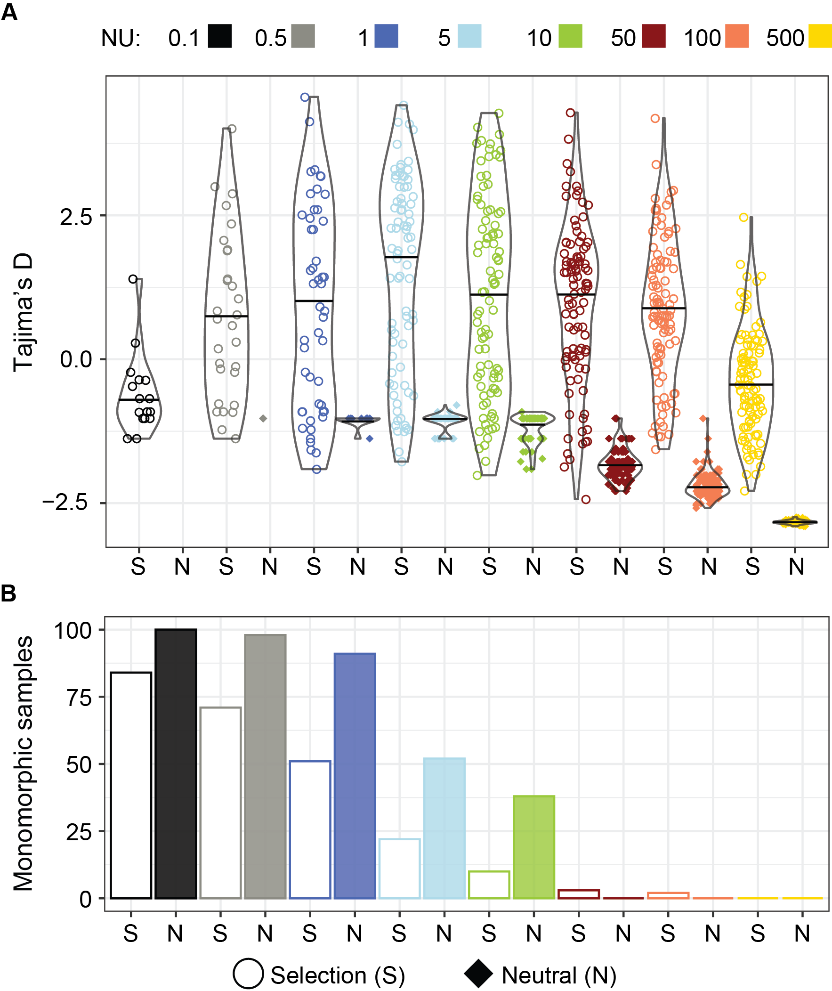

**Figure S10. Tajima’s D within and between populations.** The Tajima’s D statistics was computed for each population at the end of the adaptation (generation 10000) from a sample of $m=100$ genotypes. Samples under selection (empty circles) are compared with samples under neutrality (full diamonds). Each data point is an independent population. Those populations whose sample did not present polymorphism, are not represented (e.g. black diamonds are missing) because the Tajima’s D is not defined as the number of segregating sites would be zero. The corresponding count of monomorphic samples is given in the lower panel. Other parameters: $N={10}^{7},\rho=0, \sigma=0.05, R=2$. The violin plots show the distributions of the data and their median.

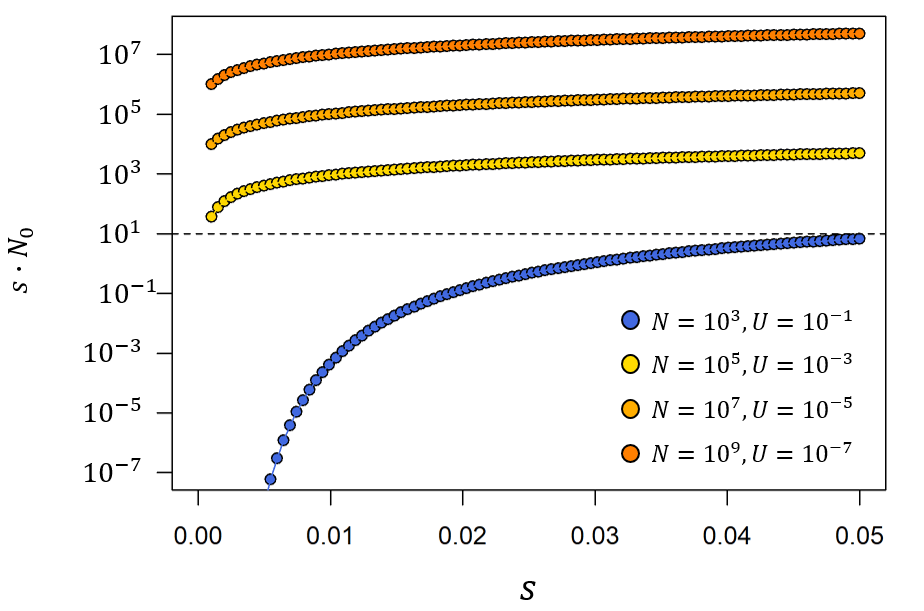

**Figure S11. From infinitesimally slow to fast Muller’s ratchet.** The rate at which the Muller’s ratchet clicks depends on the population size (N), mutation rate (U) and strength of selection (s), in a way that is not yet fully understood. The parameter $s\cdot N_{0}$, where $N_{0}:={Ne}^{-U/s}$ , was shown to be important in distinguishing different speed (Gordo and Charlesworth, 2000). Because we model mutation effects from a normal distribution (of standard deviation σ), selection $s$ is not constant as it is assumed in the classical theory of Muller’s ratchet. In order to have an approximation, we compute the $s\cdot N_{0}$ for different values of s within the interval [0.001, σ]. If $s\cdot N_{0}>10$, we expect the ratchet to click in longer times that those simulated here, while if $s\cdot N_{0}<10$ the ratchet can click. The different regimes under study have $s\cdot N_{0}\gg10$ (orange and yellow circles) except in the case with ${N=10}^{3}$ and ${U=10}^{-1}$ (blue circles) where $\sigma N_{0}=5$ and $s\cdot N_{0}$ further decreases for smaller selection. Such values of $sN_{0}$ parameter can explain why the speciation pattern observed in Fig. 6 is observable in the small populations (${N=10}^{3}$ and ${U=10}^{-1}$) but not in the large ones (${N=10}^{5},{10}^{7},{10}^{9}$).

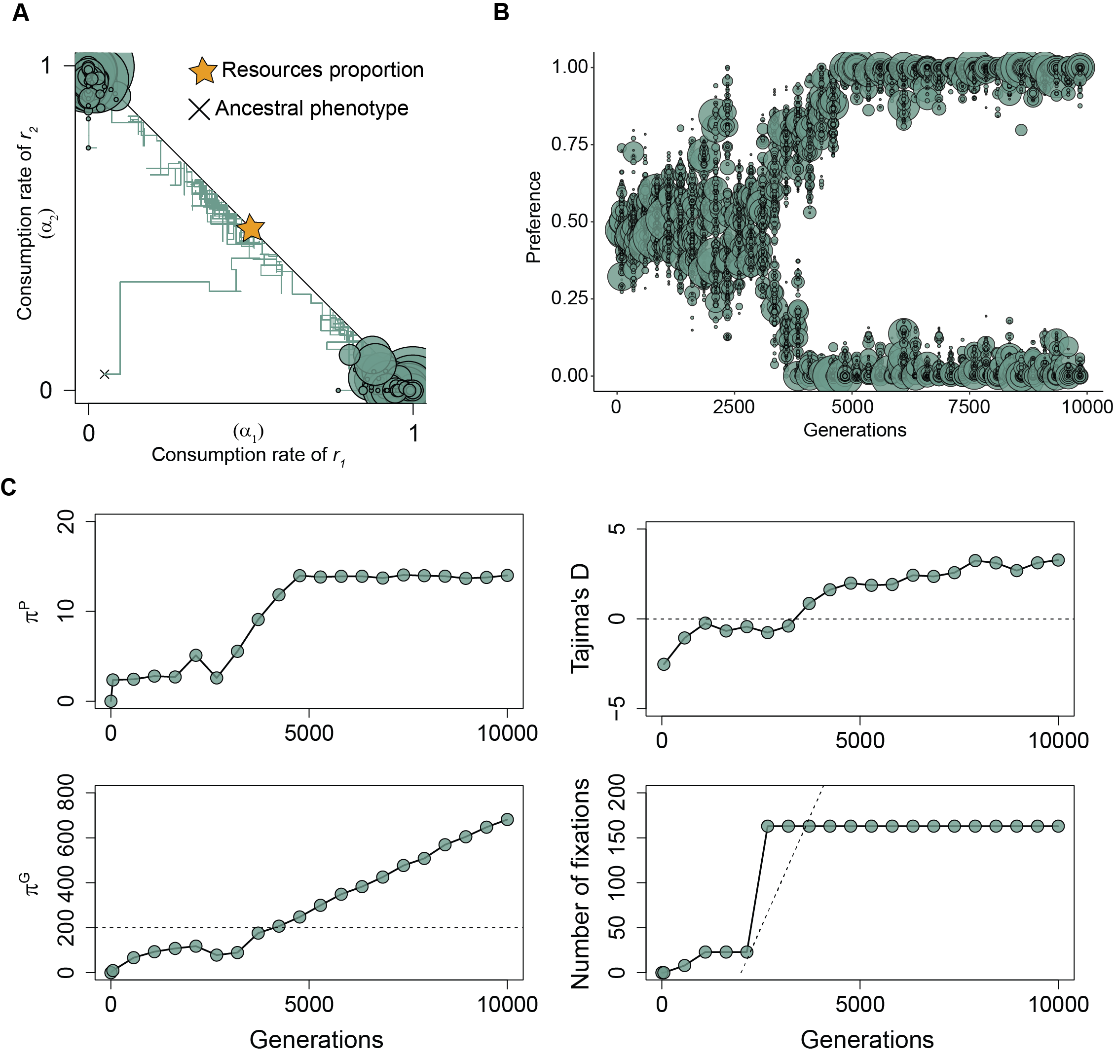

**Figure S12. Speciation in a small population with large mutational inputs.** Example population adapting with $N={10}^{3}, U={10}^{-1},\rho=0, \sigma=0.05, R=2$ for 10000 generations. A) Adaptation mapped on the phenotypic space. Each circle represents a genotype (present at generation 10000) whose size is proportional to its abundance and the lines represent the mutations. Here, diversification happens along the energetic constraint. B) Preference evolution of the population in panel A. Each circle represents a genotype (over 10000 generations) whose size is proportional to its abundance. C) $\pi_{P}$, $\pi_{G}$, Tajima’s D and number of fixations in the population from panels A-B. The dotted lines represent the expected value at neutral equilibrium. In particular $\pi_{G}=2NU$, Tajima’s D=0 and the expected rate of fixation is ∼U.

**Supplementary Tables**

|  | $a\pm2SE$ | $b\pm2SE$ |
| --- | --- | --- |
| **σ =0.05** | $-0.59\pm0.1$0 | $3.3\pm0.3$ |
| **σ =0.025** | $-0.48\pm0.09$ | $5.6\pm0.4$ |
| **σ =0.0125** | $-0.39\pm0.10$ | $8.5\pm1.0$ |

**Table S1. Diversification probability (P) increases as a logistic function of** $\mathbf{ln}\left( \mathbf{NU} \right)$ **whose shape depends on** $\boldsymbol{\sigma}$**.** Parameters of the fit the shown in the left panel of Fig. 4C. We applied the logistic function: $P=c+\frac{d-c}{1+e^{a(ln\left( NU \right)-b)}}$, where we bounded the probability $P$ between 0 and 1 (thus, $c=0, d=1$). $-a$ represents the slope of $P$ around the inflection point and $b$, the half-velocity constant ($P=0.5$ when $ln(NU)=b$). Other parameters: $NU:\left\{ 0.1, 0.2, 0.3, 0.4, 0.5, 0.6, 1, 5,10, 50,100, 500 \right\}$, $\rho=0$.

|  | $a\pm2SE$ | $b\pm2SE$ |
| --- | --- | --- |
| $\boldsymbol{\rho}$ **=0** | $-0.59\pm0.10$ | $3.3\pm0.3$ |
| $\boldsymbol{\rho}$ **=0.5** | $-0.47\pm0.07$ | $5.3\pm0.4$ |
| $\boldsymbol{\rho}$ **=1** | $-0.26\pm0.07$ | $9.5\pm1.7$ |

**Table S2. Diversification probability (P) increases as a logistic function of** $\mathbf{ln}\left( \mathbf{NU} \right)$ **whose shape depends on** $\boldsymbol{\rho}$**.** Parameters of the fit the shown in the right panel of Fig. 4C. We applied the logistic function: $P=c+\frac{d-c}{1+e^{a(ln\left( NU \right)-b)}}$, where we bounded the probability $P$ between 0 and 1 (thus, $c=0, d=1$). $-a$ represents the slope of $P$ around the inflection point and $b$, the half-velocity constant ($P=0.5$ when $ln(NU)=b$). Other parameters: $NU:\left\{ 0.1, 0.2, 0.3, 0.4, 0.5, 0.6, 1, 5,10, 50,100, 500 \right\}$, $\sigma=0.05$.

|  | $a\pm2SE$ | $b\pm2SE$ |
| --- | --- | --- |
| $\boldsymbol{N=}\boldsymbol{10}^{\boldsymbol{3}}$ | $-1.5\pm0.5$ | $3.4\pm0.3$ |
| $\boldsymbol{N=}\boldsymbol{10}^{\boldsymbol{5}}$ | $-0.51 \pm0.15$ | $3.4\pm0.6$ |
| $\boldsymbol{N=}\boldsymbol{10}^{\boldsymbol{7}}$ | $-0.56\pm0.13$ | $3.3\pm0.4$ |
| $\boldsymbol{N=}\boldsymbol{10}^{\boldsymbol{9}}$ | $-0.60\pm0.11$ | $2.9\pm0.3$ |
| $\boldsymbol{N=}\boldsymbol{10}^{\boldsymbol{5}}\boldsymbol{,}\boldsymbol{10}^{\boldsymbol{7}}\boldsymbol{or}\boldsymbol{10}^{\boldsymbol{9}}$ | $-0.55\pm0.08$ | $3.2 \pm0.2$ |

**Table S3. Diversification probability (P) increases as a logistic function of** $\mathbf{ln}\left( \mathbf{NU} \right)$ **whose shape depends on** $\boldsymbol{N}$**.** Parameters of the fit to the data shown in Fig. 6A. We applied the logistic function: $P=c+\frac{d-c}{1+e^{a(ln\left( NU \right)-b)}}$, where we bounded the probability $P$ between 0 and 1 (thus, $c=0, d=1$). $-a$ represents the slope of $P$ around the inflection point and $b$, the half-velocity constant ($P=0.5$ when $ln(NU)=b$). Other parameters: $\sigma=0.05, \rho=0$.

**Supplementary Videos**

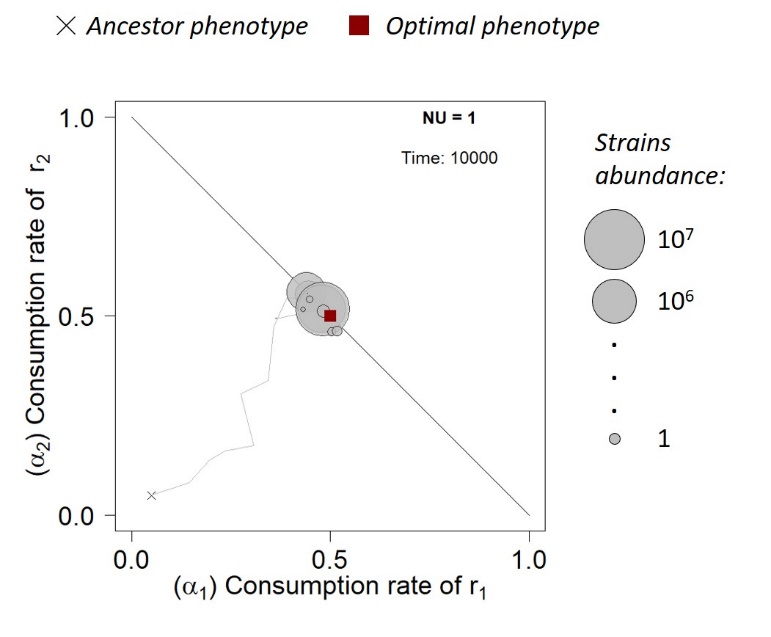

**Video S1. Evolution of a generalist phenotype.** Example population adapting with parameters $R=2, \sigma= 0.05, \rho=0.5, N={10}^{7}$and U$={10}^{-7}$. Each circle represents a genotype/phenotype whose size is proportional to its abundance in the population. The lines connecting the dots represent the effect of mutations and are shown only for the strains that are present at any given time. Time is accelerated at the end as the adaptation process does so. In this example the population adapts, via a series of selective sweeps, towards the optimal and without diversifying. The optimal phenotype (red square) corresponds to have the consumption rates mirroring the proportion of the supplied resources (50:50 in this case). To see the video, download the file “EVOLUTION_AmiconeGordo_SV1.mp4”.

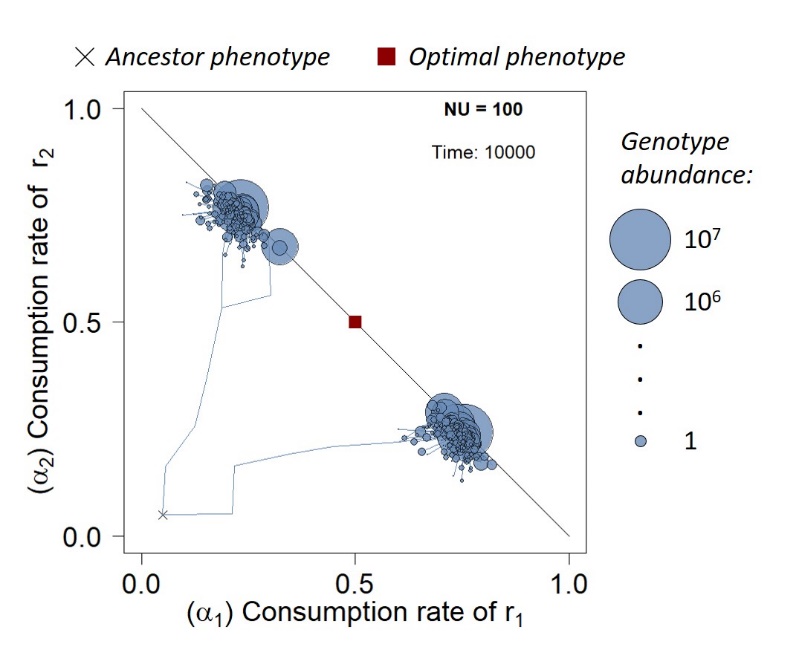

**Video S2. Ecotypes’ diversification.** Example population adapting with parameters $R=2, \sigma= 0.05, \rho=0.5, N={10}^{7}$and U$={10}^{-5}$. Each circle represents a genotype/phenotype whose size is proportional to its abundance in the population. The lines connecting the dots represent the effect of mutations and are shown only for the strains that are present at any given time. Time is accelerated at the end as the adaptation process does so. In this example the population adapts, via accumulating mutations into different lineages, towards the energy constraint but instead of reaching the optimal, it stabilizes as polymorphism. The optimal phenotype (red square) corresponds to have the consumption rates mirroring the proportion of the supplied resources (50:50 in this case). To see the video, download the file “EVOLUTION _AmiconeGordo_SV2.mp4”.
